# Long-read RNA sequencing identifies polyadenylation elongation and differential transcript usage of host transcripts during SARS-CoV-2 *in vitro* infection

**DOI:** 10.1101/2021.12.14.472725

**Authors:** Jessie J.-Y. Chang, Josie Gleeson, Daniel Rawlinson, Miranda E. Pitt, Ricardo De Paoli-Iseppi, Chenxi Zhou, Francesca L. Mordant, Sarah L. Londrigan, Michael B. Clark, Kanta Subbarao, Timothy P. Stinear, Lachlan J.M. Coin

## Abstract

Better methods to interrogate host-pathogen interactions during Severe Acute Respiratory Syndrome Coronavirus 2 (SARS-CoV-2) infections are imperative to help understand and prevent this disease. Here we implemented RNA-sequencing (RNA-seq) combined with the Oxford Nanopore Technologies (ONT) long-reads to measure differential host gene expression, transcript polyadenylation and isoform usage within various epithelial cell lines permissive and non-permissive for SARS-CoV-2 infection. SARS-CoV-2-infected and mock-infected Vero (African green monkey kidney epithelial cells), Calu-3 (human lung adenocarcinoma epithelial cells), Caco-2 (human colorectal adenocarcinoma epithelial cells) and A549 (human lung carcinoma epithelial cells) were analysed over time (0, 2, 24, 48 hours). Differential polyadenylation was found to occur in both infected Calu-3 and Vero cells during a late time point (48 hpi), with Gene Ontology (GO) terms such as viral transcription and translation shown to be significantly enriched in Calu-3 data. Poly(A) tails showed increased lengths in the majority of the differentially polyadenylated transcripts in Calu-3 and Vero cell lines (up to ~136 nt in mean poly(A) length, padj = 0.029). Of these genes, ribosomal protein genes such as *RPS4X* and *RPS6* also showed downregulation in expression levels, suggesting the importance of ribosomal protein genes during infection. Furthermore, differential transcript usage was identified in Caco-2, Calu-3 and Vero cells, including transcripts of genes such as *GSDMB* and *KPNA2*, which have previously been implicated in SARS-CoV-2 infections. Overall, these results highlight the potential role of differential polyadenylation and transcript usage in host immune response or viral manipulation of host mechanisms during infection, and therefore, showcase the value of long-read sequencing in identifying less-explored host responses to disease.

## Introduction

The Severe Acute Respiratory Corona Virus 2 (SARS-CoV-2) was first discovered in Wuhan, China at the end of 2019 and is the causative agent of the global Coronavirus Disease 2019 (COVID-19) pandemic. The World Health Organisation (WHO) reported over 5.1 million deaths and over 258 million confirmed cases globally as of late November 2021, and the global health, social and economic burden due to this disease continues to grow. Extensive research on this virus has been carried out since the first discovery of the pathogen. Nevertheless, continued exploration of the host response during an infection with SARS-CoV-2 is imperative for developing novel therapeutics, diagnostics, and prophylactics.

The host response to SARS-CoV-2 infection has been comprehensively studied within the past two years. This includes transcriptomic studies of the host using RNA sequencing (RNA-seq) from *in vitro* infections of cell lines/primary cells, *in vivo* infection models in ferrets as well as clinical samples from infected patients (Blanco-Melo et al., 2020; Mick et al., 2020; Wu et al., 2020). Of these, *in vitro* SARS-CoV-2 infection studies using continuous cell lines have been commonly used, due to the simplicity of the model. Vero (African green monkey kidney epithelial) cells are known for their high susceptibility to SARS-CoV-2, due to their defective interferon I responses (Emeny & Morgan, 1979). However, due to the lack of biological relevance using these cells, human epithelial cells have mostly been used for assessing host responses instead of Vero cells, such as Calu-3 (human lung adenocarcinoma epithelial), Caco-2 (human colorectal adenocarcinoma epithelial) and A549 (human lung carcinoma epithelial) cells. SARS-CoV-2-infected Calu-3 cells exhibited upregulation of genes involved in innate immune response to viral infections such as *IFIT2*, *OAS2*, or *IFNB1*, similar to the responses elicited by the SARS-CoV-1 virus (Blanco-Melo et al., 2020; Wyler et al., 2021). Also, in both Calu-3 and Caco-2 cells, genes involved in response to Endoplasmic Reticulum (ER) stress and mitogen-activated protein (MAP) kinases were upregulated during infection (Wyler et al., 2021). However, responses between Calu-3 and Caco-2 were found to be cell-specific. Caco-2 cells lacked in innate immune responses when infected with SARS-CoV-1/2 (Chen et al., 2021; Shuai et al., 2020; Wyler et al., 2021), and have shown fewer changes at the gene (Wyler et al., 2021) and protein level (Saccon et al., 2021) compared to Calu-3 cells. Furthermore, A549 cells have shown lack of susceptibility to SARS-CoV-2, despite being a human airway epithelial cell line like Calu-3 cells (Blanco-Melo et al., 2020; Harcourt et al., 2020). This has been attributed to the lack of the main entry receptor of SARS-CoV-2 - Angiotensin-Converting Enzyme 2 (*ACE2*) – on the surface of these cells. However, air-liquid interface culturing or *ACE2*-expressing A549 (A549-hACE2) cells enhanced the susceptibility to SARS-CoV-2 (Sasaki et al., 2021; Xie et al., 2020). Overall, host responses appeared to vary between different epithelial cell lines and were dependent on the multiplicity of infection (MOI) of the virus in A549-h*ACE2* cells (Blanco-Melo et al., 2020).

Most RNA-seq data reported in the literature have been generated using short-read sequencing methods such as Illumina sequencing (Bibert et al., 2021; Islam et al., 2021). In these studies, differential expression and Gene Ontology (GO)/ Kyoto Encyclopedia of Genes and Genomes (KEGG) pathway analyses have been the main outcomes. Short-read RNA-seq is an effective technique for measuring differential mRNA abundance. However, utilising a long-read sequencing platform provides the ability to discern other functionally significant mRNA features such as length of the poly(A) tails, alternative splicing, and differential isoform usage (de Jong et al., 2017; De Paoli-Iseppi, Gleeson, & Clark, 2021; Gleeson et al., 2021; Workman et al., 2019). These additional mRNA features have been linked with different disease states (Curinha, Oliveira Braz, Pereira-Castro, Cruz, & Moreira, 2014; Dick et al., 2020; Tazi, Bakkour, & Stamm, 2009). However, these events have not been studied in depth for infectious diseases, especially with SARS-CoV-2 infections. An ability to measure full-length transcripts, polyadenylation status and isoform usage would permit significantly enriched insights into host responses to viral infection than standard RNA-seq methods allow.

Here we report the use of RNA-seq methods from the Oxford Nanopore Technologies (ONT) platform (direct RNA, direct cDNA and PCR cDNA) to carry out an in-depth investigation into the host response to SARS-CoV-2 *in vitro*. The responses were visualised throughout a time-course (0, 2, 24 and 48 hours post infection (hpi)) using four epithelial cell lines (Vero, Calu-3, Caco-2 and A549). Previously we performed a comprehensive analysis of the viral response for some of these datasets (Chang et al., 2021). In this current study, we investigated differential polyadenylation and transcript usage between infected and mock control cells. Additionally, we were interested in whether long-read differential expression analysis conveyed similar differential expression results to short-read RNA-seq studies shown in literature. Overall, our study demonstrated the value of long-read sequencing in identifying less-explored host responses to disease.

## Methods

### Data availability

ONT sequencing data (direct RNA and direct cDNA) for this study from cell lines (Vero, Caco-2 and Calu-3) was derived from our previous work (Chang et al., 2021), and is currently publicly available at NCBI repository BioProject PRJNA675370. Additional datasets were generated for this study including PCR cDNA datasets for cell lines (Vero, Caco-2, Calu-3 and A549) and the direct RNA and direct cDNA datasets for A549. These datasets are also available at NCBI repository BioProject PRJNA675370. The results of individual analyses are available at Figtree DOI: <u>10.6084/m9.figshare.17139995</u> (differential expression), <u>10.6084/m9.figshare.16841794</u> (differential polyadenylation) and <u>10.6084/m9.figshare.17140007</u> (differential transcript usage).

### Experimental methods

#### Cell culture and RNA extraction/preparation

Cell culture and RNA extraction/preparation methods have been described previously (Chang et al., 2021) for Calu-3 (human lung adenocarcinoma epithelial - ATCC HTB-55), Caco-2 (human colorectal adenocarcinoma epithelial - ATCC HTB-37) and Vero (African green monkey kidney epithelial - ATCC CCL-81) cells. For this current study, we additionally cultured A549 (human lung carcinoma epithelial – ATCC CCL-185) cells to supplement our main data, using similar methods. Briefly, A549, Vero, Calu-3 and Caco-2 cell lines were cultured in T75 flasks and maintained at 37 °C and 5% (v/v) CO_2_. A549 cells were cultured with Ham’s F-12K (Kaighn’s) Medium (Gibco) supplemented with 10% FBS, 4 mM L-glutamine (Media Preparation Unit, The Peter Doherty Institute for Infection and Immunity (Doherty Institute)), 100 IU penicillin, 10 μg streptomycin/mL, 1X non-essential amino acids (Gibco-BRL) and 50 μM B-mercaptoethanol (Life Technologies). All cell lines were seeded in 4 x 6-well tissue-culture plates and maintained at 70-80% confluency for infection. Three wells of the 6-well plates were infected with SARS-CoV-2 (Australia/VIC01/2020) at a MOI of 0.1 and the remaining wells were used as mock controls for four time points (0, 2, 24 and 48 hpi). Total cellular RNA was extracted with the RNeasy Mini Kit (Qiagen), treated with the Turbo DNAse-free Kit (Invitrogen) and purified with RNAClean XP magnetic beads (Beckman Coulter). The final resulting RNA was eluted in nuclease-free water. Quality control was carried out using NanoDrop 2000C (Thermo Fisher Scientific), Bioanalyzer 2100 (Agilent Technologies) and Qubit 4 Fluorometer (Invitrogen).

#### Library preparation and sequencing

Library preparation and sequencing methods have been described previously (Chang et al., 2021). Briefly, RNA from mock control and infected cells harvested at 0, 2, 24 and 48 hpi from Caco-2, Calu-3 and Vero cells was sequenced with the ONT Direct cDNA Sequencing Kit (SQK-DCS109) in conjunction with the Native Barcoding Kit (EXP-NBD104). RNA harvested at 2, 24 and 48 hpi was sequenced with the Direct RNA Sequencing Kit (SQK-RNA002) by pooling the RNA from replicate wells. For this current study, RNA from A549 cells was sequenced as per our previous work with minor modifications in the number of time points sequenced, to supplement our main data. The ONT Direct RNA Sequencing Kit (SQK-RNA002) was used to prepare 6 μg of pooled total RNA (2 μg RNA from each replicate well) from control and infected cells at the 24 hpi. The Direct cDNA Sequencing Kit (SQK-DCS109) was used in conjunction with the Native Barcoding Kit (EXP-NBD104) to prepare 3 μg of total RNA from all control and infected replicates separately at both 0 and 24 hpi time points. All direct RNA and direct cDNA libraries were loaded onto a R9.4.1 MinION flow cell and sequenced for 72 hrs using an ONT MinION or GridION. Additionally, PCR cDNA long-read sequencing was carried out with RNA from all four cell lines (Vero, Caco-2, Calu-3 and A549 cells) using the following methods: cDNA libraries were constructed with the PCR-cDNA Sequencing (SQK-PCS109) and PCR Barcoding (SQK-PBK004) kits using the supplied protocol. RNA samples from 0 and 24 (hpi) were randomised and multiplexed for sequencing in groups of six using sequential barcodes. 100 ng of sample RNA was used for cDNA synthesis. Transcripts were amplified by PCR and barcodes added using the specified cycling conditions with a 7 min extension time and 13x cycles. Amplified samples were individually cleaned using 0.5x AMPure XP beads (Beckman Coulter) and quantified using a Qubit 4 Fluorometer (Invitrogen). The length distribution was determined via the TapeStation 4200 (Agilent Technologies) before pooling. Equimolar amounts of each barcoded sample were pooled to a total of 100 – 200 fmol (assuming median transcript size = 1.1 kb). 100 fmol of final libraries were loaded onto a R9.4.1 MinION flow cell and sequenced for 72 hrs on an ONT GridION. Run metrics were monitored live and if active pores dropped below 200, any remaining library was loaded following a nuclease flush. Synthetic ‘sequin’ RNA standards, provided in two mixes (A and B) (Hardwick et al., 2016), were added to each sample in direct RNA and PCR cDNA libraries. Mix A and B sequins, diluted 1:250 (approximately 6-10% of estimated total mRNA), were added to infected and control samples, respectively.

#### Data analysis

Publicly available data from our previous work (Chang et al., 2021) in combination with data generated from this study were analysed using Spartan (Meade, Lafayette, Sauter, & Tosello, 2017) and Nectar from the Australia Research Data Commons.

#### Basecalling, alignment and generating counts files

All FAST5 files were basecalled using standalone *Guppy* v3.5.2 (https://community.nanoporetech.com/sso/login?next_url=%2Fdownloads), except PCR cDNA data from Vero cells which were live-basecalled using *Guppy* v3.2.8. All resulting FASTQ data were mapped using *Minimap2* v2.17 (Heng Li, 2018). Direct RNA-seq data was mapped to the combined genome (consisting of human/African green monkey genome from Ensembl (release 100), SARS-CoV-2 Australia virus (Australia/VIC01/2020, NCBI:MT007544.1) and the RNA sequin decoy chromosome genome (Hardwick et al., 2016) with the default direct RNA parameters ‘-ax splice -uf -k14 --secondary=no’ and for all cDNA datasets ‘-ax splice –secondary=no’. All data were mapped to the respective combined transcriptome using the following parameters – ‘-ax map-ont’. The resulting BAM files were sorted and indexed using *Samtools* v1.9 (H. Li et al., 2009). Counts files were generated using *Featurecounts* v2.0.0 (Liao, Smyth, & Shi, 2014) for genome-mapped cDNA data, and with *Salmon* v0.13.1 (Patro, Duggal, Love, Irizarry, & Kingsford, 2017) for transcriptome-mapped cDNA data.

#### Differential expression analysis

*DESeq2* was used to identify differentially expressed genes/transcripts from direct cDNA data. A minimum expression threshold of five reads per gene/transcript across all the replicates was used. Comparisons between control and infected cells were made per time point (0, 2, 24, 48 hpi) with standard methods. Also, the changes between time points (0-2, 0-24, 2-24, 24-48 hpi) were compared, where the interaction term between control and infected across time points were found using a method by Steven Ge (https://rstudio-pubs-static.s3.amazonaws.com/329027_593046fb6d7a427da6b2c538caf601e1.html#example-4-two-conditionss-three-genotpes-with-interaction-terms). This is a more sensitive method as it calculates the changes between time points in infected cells while accounting for the changes in the expression level in the control cells. All genes/transcripts with padj < 0.05 were regarded as significantly differentially expressed. The heatmap of differentially expressed genes in Caco-2 and Calu-3 at 24 and 48 hpi were generated using *multiGO*. The filters utilised were padj < 0.05, GO p-value < 0.0001, scaled by row, with ten maximum GO terms. Columns with more than 80% of NA’s and rows with more than 10% of NA’s were excluded.

#### Poly(A) tail length analysis

Two tools were used for poly(A) tail length analysis; *nanopolish* (Simpson et al., 2017) and *tailfindr* (Krause et al., 2019). For the *nanopolish* analysis, all Caco-2, Calu-3 and Vero direct RNA BAM files mapped to the combined reference genome (host, sequin, virus) were indexed with the *nanopolish* v0.13.2 ‘index’ function with the command ‘nanopolish index -d $FAST5 -s $SEQUENCING_SUMMARY $FASTQ’. The poly(A) tail lengths of each read were estimated using the ‘polya’ function with default parameters ‘nanopolish polya --reads $FASTQ --bam $SORTED_BAM --genome $COMBINED_REFERENCE_GENOME > output.tsv’. The host reference sequence names were extracted from the reference file with the following command ‘cat $REFERENCE_GENOME | grep ‘>’ | cut -d ‘’ -f 1 | cut -f 2 -d ‘>’’. Using this name file, the host reads were extracted from the final *nanopolish* TSV file by this command ‘awk ‘NR==FNR{A[$1]; next} $2 in A’$NAMES.TSV $TSV’.

After the poly(A) lengths were determined, duplicates were removed, and the outputs were merged with a file generated by an in-house pipeline – *npTranscript* - which allowed read names to be associated with Ensembl ID’s. The data were grouped per Ensembl ID and whether they were mitochondrial or non-mitochondrial genes, and the median poly(A) lengths were calculated. Differential polyadenylation between the overall median lengths of control vs infected cells per cell line were determined with p-values using Wilcoxon’s test of ranks for Ensembl ID’s with more than one entry.

As the *nanopolish* results revealed evidence of differential polyadenylation between the overall median of control and infected poly(A) lengths, *tailfindr* analysis was utilised to gather more evidence at a gene level. For *tailfindr* analysis, direct cDNA datasets from Vero, Calu-3 and Caco-2 were utilised. Basecalled FAST5 files were subsetted by read ID’s derived from demultiplexed FASTQ files. Each replicate was passed through *tailfindr* v0.1.0 separately. The median poly(A) and poly(T) lengths were calculated per gene and grouped by whether they were mitochondrial or non-mitochondrial.

The Pearson product-moment correlations between the median poly(A) and poly(T) lengths per gene from *tailfindr* analyses were compared for 2, 24 and 48 hpi datasets from Caco-2, Calu-3 and Vero datasets. Additionally, the Spearman’s correlations between the *tailfindr* poly(T)/(A) and *nanopolish* poly(A) median lengths per gene (in control and infected cells) were compared for Calu-3 48 hpi datasets via the ‘cor.test’ function in the *stats* package in R.

*tailfindr* poly(T) results were used for the main polyadenylation linear mixed-model analysis as replicate information was able to be preserved and showed higher correlation to *nanopolish* poly(A) lengths compared with *tailfindr* poly(A) lengths. The raw poly(T) lengths were log-transformed due to the right-skew distribution and data with at least 6 entries were selected. Then, the package *lmerTest* v3.1-3 (Kuznetsova, Brockhoff, & Christensen, 2017) was used to derive a linear mixed-effects regression (lmer) and therefore calculate the effect of SARS-CoV-2 infection compared with control mock-infected cells. The p-values were generated per gene and Benjamini-Hochberg adjusted using the ‘p.adjust’ function in R, which were filtered by padj < 0.05. Raincloud plots were generated for median poly(T) lengths of each gene with increased poly(A) length in the Calu-3 48 hpi dataset in both conditions (control and infected) using using *ggplot2* v3.3.4 (Wickham, 2016) to replicate the raincloud plots generated by the *raincloudplots* package in R (Allen et al., 2021).

To test whether the same significant genes in the mixed-model analysis appeared in *nanopolish* Calu-3 48 hpi poly(A) data, the raw tail lengths were log-transformed and the median lengths per gene were tested between control and infected cells using Wilcoxon’s test of ranks, where p-values were adjusted using Benjamini-Hochberg adjustment as above.

#### Differential transcript usage analysis

Counts from *Salmon* using transcriptome-mapped BAM files were used to determine the differential transcript usage of transcripts between control and infected conditions for each cell line and time point. The counts were input into *DRIMSeq* v1.16.1 (Nowicka & Robinson, 2016) and filtered by conditions (min_samps_gene_expr = 6, min_samps_feature_expr = 3, min_gene_expr = 10, min_feature_expr = 10). The output was used for stage-wise analysis using *StageR* v1.10.0 (Van Den Berge, Soneson, Robinson, & Clement, 2017), where the final list of significant genes and transcripts was filtered by padj < 0.05.

#### GO and KEGG pathway analysis

Significant biological GO biological terms and KEGG pathways were identified with genes that were found to be significantly differentially expressed and polyadenylated in the analyses above. For differential expression analysis, genes found to be differentially expressed in direct cDNA datasets for each condition and time point were used for analysis. For differential polyadenylation analysis, genes that were found to be increased and decreased in poly(A) length in Calu-3 48 hpi direct cDNA dataset were used for analysis. All pathway analyses were carried out using a novel shiny-app *multiGO* (http://coinlab.mdhs.unimelb.edu.au/multigo). *multiGO* uses a hypergeometric test against a background of all genes included in the *GO annotation database* v100. For differential expression, thresholds of padj < 0.05 and enrichment p-value < 1E-6 in at least one dataset were used for generating the GO plot, and thresholds of padj < 0.05 and enrichment p-value < 0.0001 were used for generating the KEGG plot. Non-significant bubbles were also shown (http://coinlab.mdhs.unimelb.edu.au/multigo/?subdir=multigo/multiGO&file=DESeq2.zip). All terms with padj < 0.05, enrichment p-value < 0.05 and at least two genes were deemed as significant for the analysis. For differential polyadenylation and differential polyadenylation vs expression analyses, thresholds of padj < 0.05 and enrichment p-value < 0.0001 were used (http://coinlab.mdhs.unimelb.edu.au/multigo/?subdir=multigo/multiGO&file=DP_48hpi_no_filter_merged.zip).

#### Differential expression vs differential polyadenylation

Using a hypergeometric test, the probability of obtaining greater than or equal to two genes overlapping between the differential expression and polyadenylation analyses were tested. Counts of downregulated genes (padj < 0.05) from Calu-3 48 hpi datasets from *DESeq2* analysis were set as *m*=253. Counts of genes with elongated poly(A) tails were set as *k*=13. Genes which were both downregulated and increased in poly(A) length were set as *n*=2. The total background count was set as N=15,426. The code used for the probability calculation was ‘phyper(x,k,15426-k,m,lower.tail=F) + dhyper(x,k,15426-k,m)’ and was carried out in R. Both mitochondrial and non-mitochondrial genes were included in this calculation.

## Results

### Viral burden changes between different cell lines and over time

We utilised the percentage of mapped reads to host or virus to assess the level of infection in each cell line across different datasets and over time (**Data S1**) to add to our earlier study (Chang et al., 2021). Between 0 and 2 hpi, the percentage of viral reads were minimal (< 0.1% of all reads) across each of the cell lines. At 24 hpi, differences between the cell lines started to appear, with Vero cells leading in infection with ~45% of reads mapping to virus, followed by Caco-2 (~2.3%), Calu-3 (~2%), and A549 (< 0.01%), as measured with direct cDNA datasets (**Data S1**). The relative proportions of viral transcripts between these four cell lines were aligned with the results in literature at the 24 hpi (Saccon et al., 2021). The final time point (48 hpi) showed the greatest per-cell-line infection in Caco-2 (~12.5%) and Calu-3 (~3.7%) cells but lowered in percentage in Vero cells (~25%) compared with 24 hpi. These results agreed with the idea that the infection peaked at 24 hpi in Vero cells as shown by our previous study (Chang et al., 2021). Interestingly, the percentage of reads mapping to virus were markedly higher in direct RNA datasets compared with the direct cDNA and PCR cDNA datasets at the 24 hpi (**Table 1**). The reason for this may be due to the direct RNA method involving the sequencing of the mRNA molecule, instead of the reverse-transcribed cDNA, as in the direct cDNA and PCR cDNA methods. This would remove any biases caused by the reverse-transcription. These results suggested that measuring viral infection using more than one ONT RNA-seq approach may be more beneficial to accurately gauge the level of viral RNA in the sample.

**Table 1.** Proportions of average viral reads in 24 hpi datasets in Vero, Calu-3, Caco-2 and A549 cell lines. Related to **Data S1**.

| Cell line | Direct RNA (%) | Direct cDNA (%) | PCR cDNA (%) |
| --- | --- | --- | --- |
| Vero | 74 | 45 | 55 |
| Calu-3 | 4 | 2 | 3 |
| Caco-2 | 4 | 2 | 3 |
| A549 | 0.02 | 0.01 | 0.01 |

### Cell-type specific changes in host gene expression *in vitro* following virus infection using long-read sequencing

The host responses to SARS-CoV-2 have been extensively studied at the gene and protein expression level (Saccon et al., 2021; Shah, Firmal, Alam, Ganguly, & Chattopadhyay, 2020). As long-read sequencing enables full-length transcripts to be sequenced unlike short-read sequencing, we were interested in whether our long-read differential expression results would reveal similar results to existing studies (Blanco-Melo et al., 2020; Chen et al., 2021; Saccon et al., 2021; Sun et al., 2021). The direct cDNA datasets were used for differential expression analysis as it included data from all four time points (0, 2, 24, 48 hpi) in Calu-3, Caco-2 and Vero cells, and two time points (0 and 24 hpi) in A549 cells.

It is well-known that A549 cells are invulnerable to SARS-CoV-2, due to the lack of *ACE2* receptors (Blanco-Melo et al., 2020). However, *ACE2* is expressed in varying degrees in different human tissues (M.-Y. Li, Li, Zhang, & Wang, 2020) and is expressed relatively poorly in the respiratory tract (M.-Y. Li et al., 2020). Previous studies have shown the expression of *ACE2* at the gene (Puray-Chavez et al., 2021) and protein (Saccon et al., 2021) level in Vero, Calu-3 and Caco-2 cells. Furthermore, the importance of the protease *TMPRSS2* during SARS-CoV-2 has been noted (Hoffmann et al., 2020). Our long-read data were in line with some of these results, where no transcripts mapped to *ACE2* and *TMPRSS2* in A549 cells. However, we observed the absence/low expression of *ACE2* (< 5 reads per replicate) and *TMPRSS2* (< 25 reads per replicate) genes across all our susceptible cell lines. The presence of these transcripts correlated with the viral burden observed in each cell line from our previous study (**Data S1**) (Chang et al., 2021).

In all cell lines, the earlier infection time points (0 and 2 hpi) showed little significant differential expression as expected, given the short period of infection in which host responses could be elicited. We observed an increase in significantly upregulated genes in Calu-3 and Vero cell lines at 24 hpi (**Figure 1**). At the final time point (48 hpi), we noted an increase in downregulated genes as well as the presence of upregulated genes in Calu-3, Vero and Caco-2 cell lines. In line with previous studies (Saccon et al., 2021; Wyler et al., 2021), while Calu-3 and Vero cells exhibited clear changes in transcriptional activity throughout the final two time points, Caco-2 cells revealed little differential expression activity (**Figure 1 & Table 2**). These results were recapitulated in a second measurement of gene expression changes (**Figure S1**). In this combined analysis, the differences between the host gene expression of control and infected cells across two time points were measured (interaction term – see **Methods**) as opposed to differences at each individual time point.

**Figure 1.**
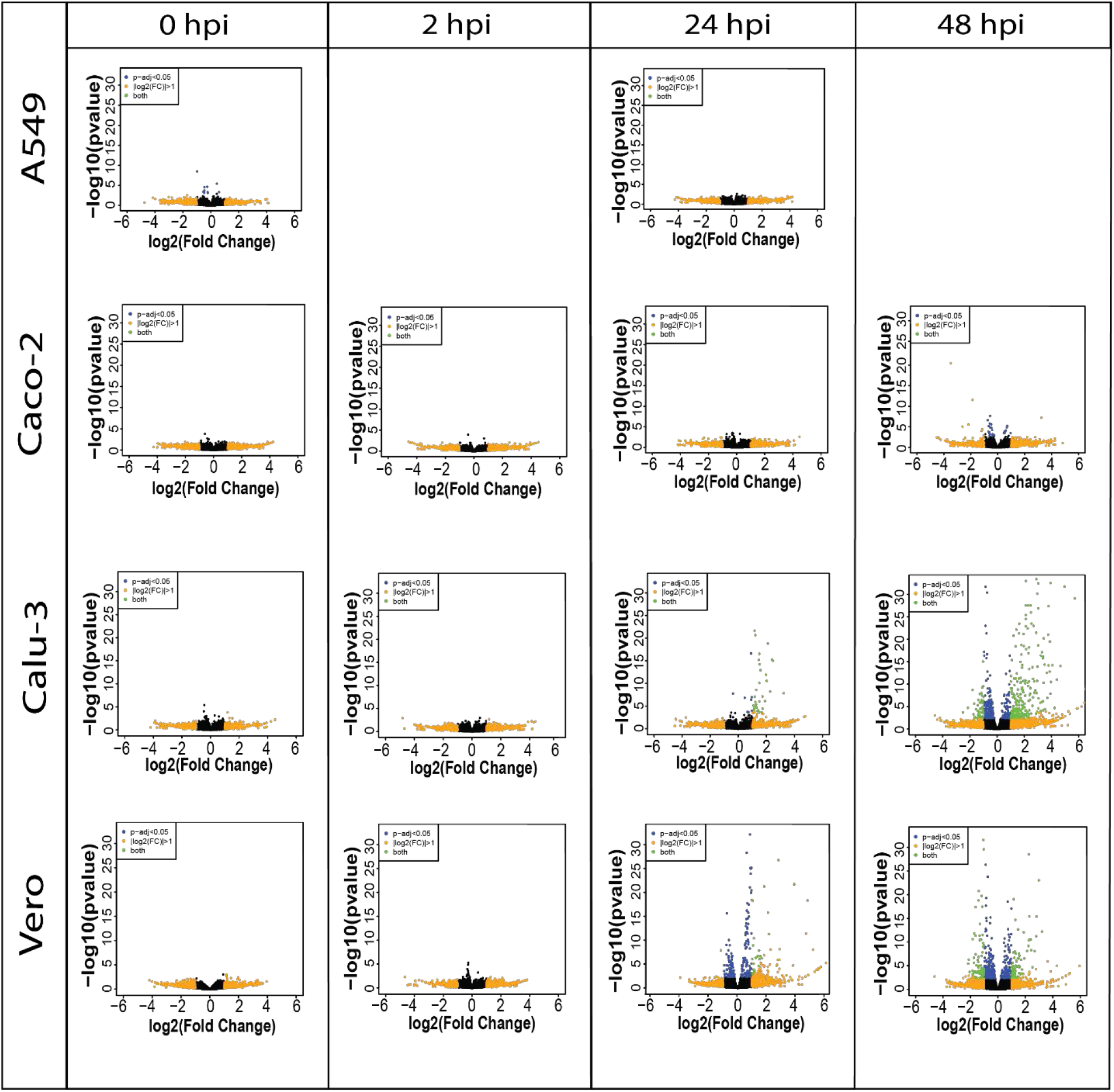
Volcano plots show the difference in expression level between control and infected cells per cell line (A549, Caco-2, Calu-3 and Vero) in direct cDNA datasets using *DESeq2*. X-axis represents log2FC and Y-axis displays −log10 p-adjusted value (padj). 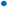 padj < 0.05 (blue), 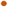 |log2FC |> 1 (orange), 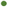 both (green). Increased number of differentially expressed genes are shown in latter time points in Caco-2, Calu-3 and Vero cell lines, and a lack of differential expression was observed in A549 cells. Related to **Figure S1 & Table 2**.

**Table 2.** The number of significantly differentially expressed genes (padj < 0.05) between control and infected cells in A549, Caco-2, Calu-3 and Vero cells over 2, 24 and 48 hpi in direct cDNA datasets. Related to **Figure 1 & S1**.

| Cell line | Time point | Up | Down |
| --- | --- | --- | --- |
| A549 | 24 | 0 | 0 |
| Caco-2 | 2 | 0 | 0 |
| Caco-2 | 24 | 0 | 0 |
| Caco-2 | 48 | 8 | 17 |
| Calu-3 | 2 | 0 | 0 |
| Calu-3 | 24 | 41 | 1 |
| Calu-3 | 48 | 371 | 253 |
| Vero | 2 | 0 | 0 |
| Vero | 24 | 134 | 111 |
| Vero | 48 | 210 | 290 |

### Calu-3 and Caco-2 cells show distinct gene expression level patterns

As the initial differential expression results showed differences between Calu-3 and Caco-2 cell lines, we then investigated the similarity of gene expression patterns between the two cell lines at the 24 and 48 hpi via a heatmap (**Figure 2**). Following the results in literature (Chen et al., 2021; Wyler et al., 2021), our results also showed higher relative expression of interferon-related genes such as *IFI6* and *IFITM3* in infected Calu-3 cells compared with Caco-2 cells. This was observed especially at the 48 hpi time point. In contrast, Caco-2 cells revealed higher expression of ribosomal protein genes as well as mitochondrial genes compared with Calu-3 cells. Therefore, our long-read results supported the idea that Caco-2 and Calu-3 cells have distinct gene expression level patterns, and that Caco-2 cells have diminished innate immune responses in contrast to Calu-3 cells.

**Figure 2.**
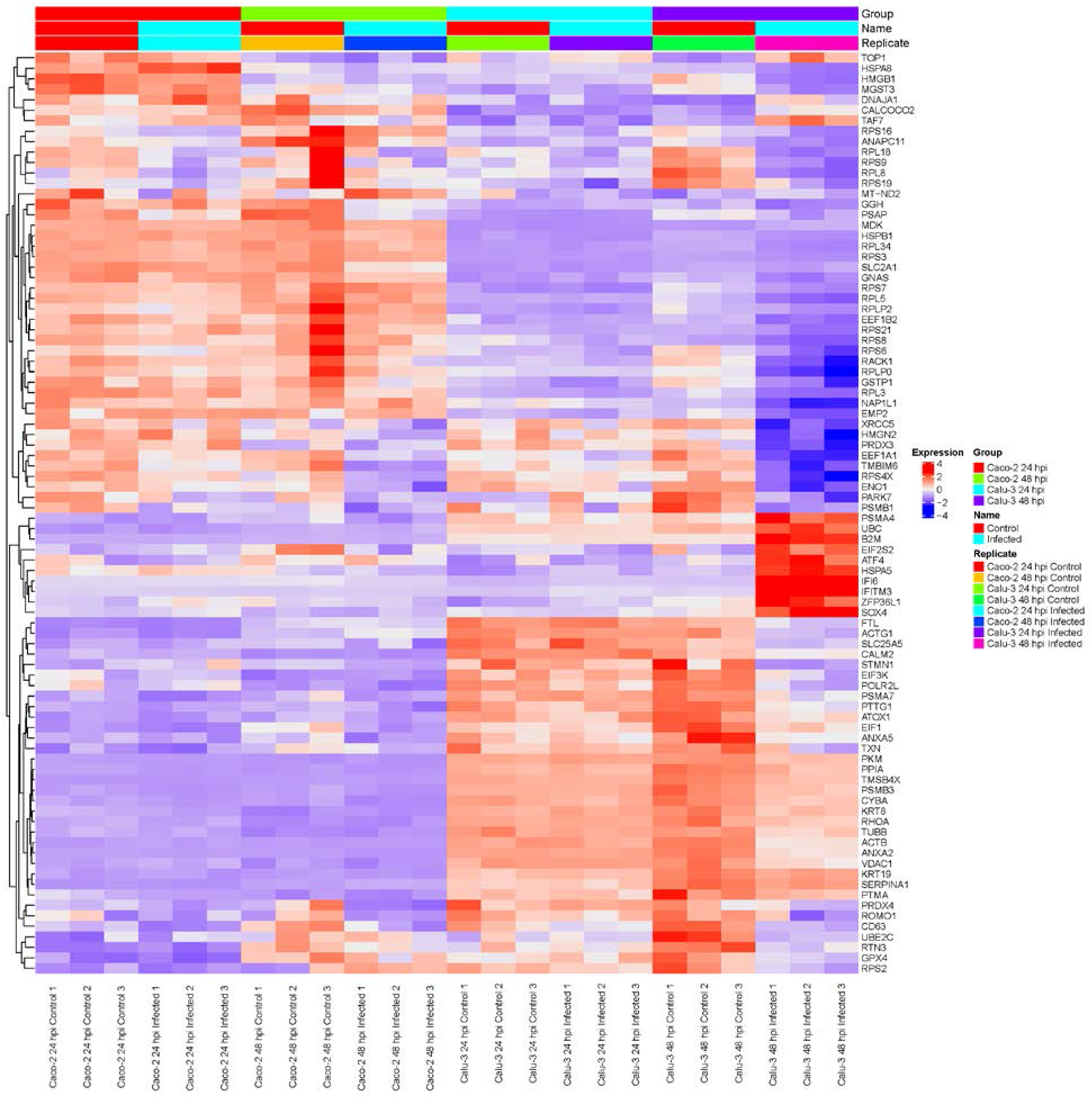
Heatmap of relative gene expression in Caco-2 and Calu-3 cells at 24 and 48 hpi using direct cDNA datasets reveal distinct gene expression profiles in each cell line, analysed by *DESeq2* and visualized by *multiGO*. The data was filtered by padj < 0.05, enrichment p-value < 0.0001. The expression levels were scaled per row and organised based on relevant GO terms. In Calu-3 cells, higher relative expression of interferon-response genes was observed compared with Caco-2 cells. In contrast, Caco-2 cells showed higher relative expression of ribosomal protein and mitochondrial genes. Related to **Figure 1 & S1 & Table 1**.

### GO and KEGG analyses reveal similarities between Calu-3, Caco-2 and Vero cells

To investigate the cell-specific gene expression changes at a deeper level, we wondered whether any enrichment of pathways was shared between multiple cell types. By utilising the genes which were significantly differentially expressed in the direct cDNA data, GO biological and KEGG pathway analyses were carried out using a new visualisation tool *multiGO* (http://coinlab.mdhs.unimelb.edu.au/multigo) (**Figures 3 & S2**).

**Figure 3.**
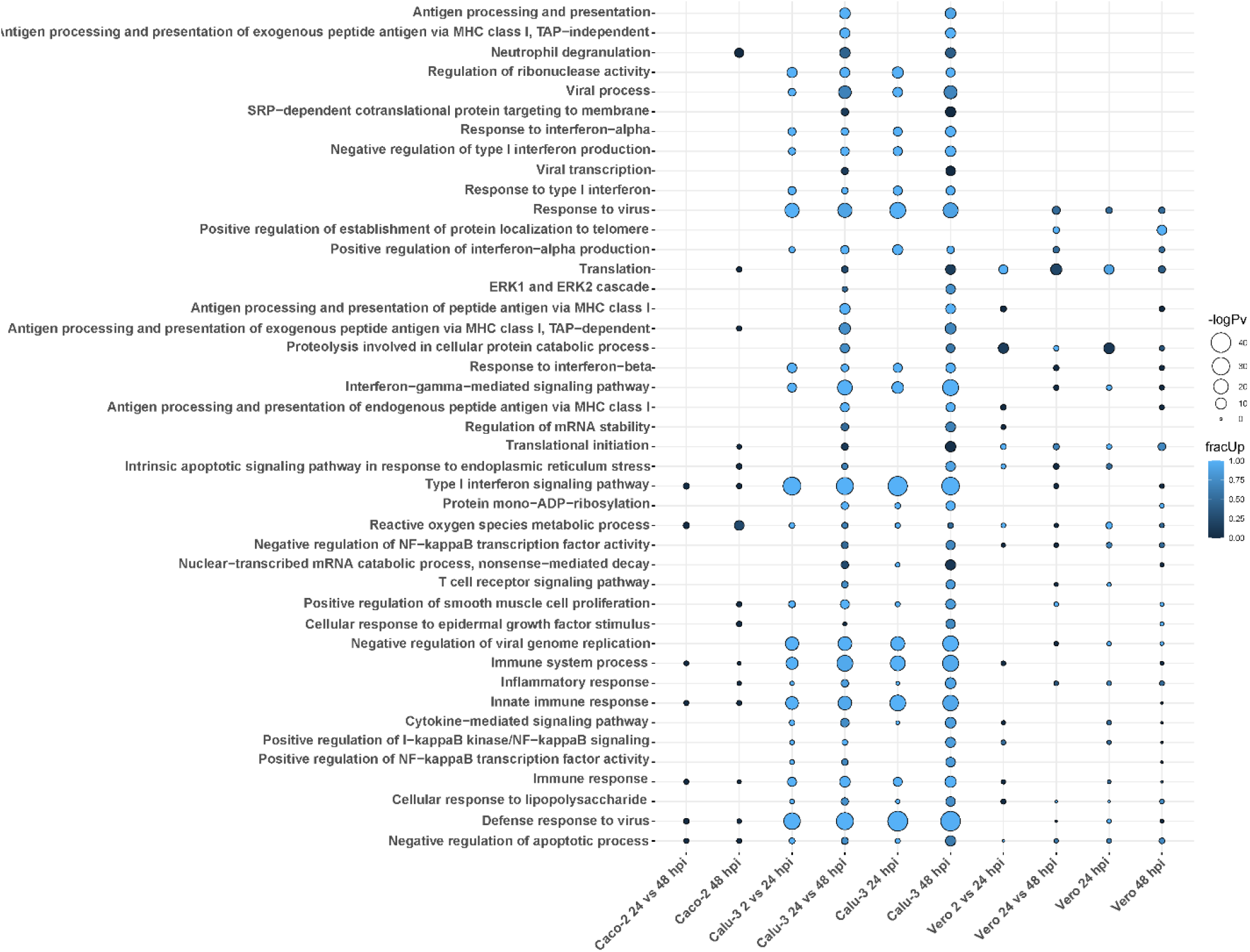
GO biological terms of differentially expressed genes in Calu-3, Caco-2 and Vero direct cDNA datasets analysed by *DESeq2* and visualised with *multiGO*. Results include datasets (in order); Caco-2 24 vs 48, Caco-2 48, Calu-3 2 vs 24, Calu-3 24 vs 48, Calu-3 24, Calu-3 48, Vero 2 vs 24, Vero 24 vs 48, Vero 24 and Vero 48 hpi. Commonly enriched GO terms across the three cell lines involved the *ROS metabolic process*. The bubble size and colour indicate the −log10 enrichment p-value and fraction of upregulated genes, respectively. Thresholds of padj < 0.05 and enrichment p-value < 1E-6 in at least one dataset were used for generating the plot, where insignificant bubbles are also shown on the plot. All terms with padj < 0.05, enrichment p-value < 0.05 involving at least two genes were deemed as significant for the analysis. Related to **Figure S2 & Data S2**.

Amongst many enriched GO biological terms, *reactive oxygen species (ROS) metabolic process* was enriched in all three SARS-CoV-2 susceptible cell lines. Also, we found that only *neutrophil degranulation* was commonly enriched exclusively in the two human cell lines and absent in Vero cells. In contrast, a greater number of terms were shared between Calu-3 and Vero cell lines. These terms included *response to virus*, *positive regulation of interferon-alpha production* and *translation* (**Figure 3 & Data S2**).

As expected, the Calu-3 cell line showed an increase in innate immune responses, with the strongest GO enrichment for various terms associated with host immune responses to pathogens. This included terms such as *defense response to virus* and *type I interferon signalling pathway* (**Figure 3 & Data S2**). As shown above with the gene expression results, these responses were either absent or lacking in Caco-2 cells compared with Calu-3 cells. Additionally, some unique GO terms were enriched in Vero cells. This included *positive regulation of establishment of protein localization to telomere* (**Figure 3 & Data S2**).

Similarly, Calu-3 cells presented with the strongest significant enrichment of KEGG pathways (**Figure S2 & Data S2**). In our data, the enriched pathways were related to viral infections such as *influenza A* (*MX1*, *OAS1*, *OAS2*, *OAS3*, *RSAD2*, *STAT1*) and *measles* (*MX1*, *OAS1*, *OAS2*, *OAS3*, *STAT1*). These pathways were upregulated at the 24 and 48 hpi time points as well as between 2 vs 24 hpi and 24 vs 48 hpi datasets in infected cells compared with control cells (**Figure S2**). *DDX58* was also observed as upregulated in these pathways in the same datasets except for 2 vs 24 hpi. This has also been shown in influenza A studies (Watson et al., 2020) and the gene has been shown to encode a cytosolic sensor for other coronaviruses ((J. Li, Liu, & Zhang, 2010). The *coronavirus disease* pathway was enriched in both Calu-3 and Vero cells (**Table 3**). Also, as shown previously in various infected epithelial cell lines (Martinelli, Akhmedov, & Kwee, 2021), pathways related to neurological diseases such as Alzheimer’s, Parkinson’s and Huntington’s diseases were found to be enriched in both Vero and Calu-3 cells. The majority of genes in these pathways were downregulated (**Figure 3 & Data S2**). Overall, our long-read RNA-seq data were aligned with results from previous studies which utilised short-read RNA-seq.

**Table 3.**
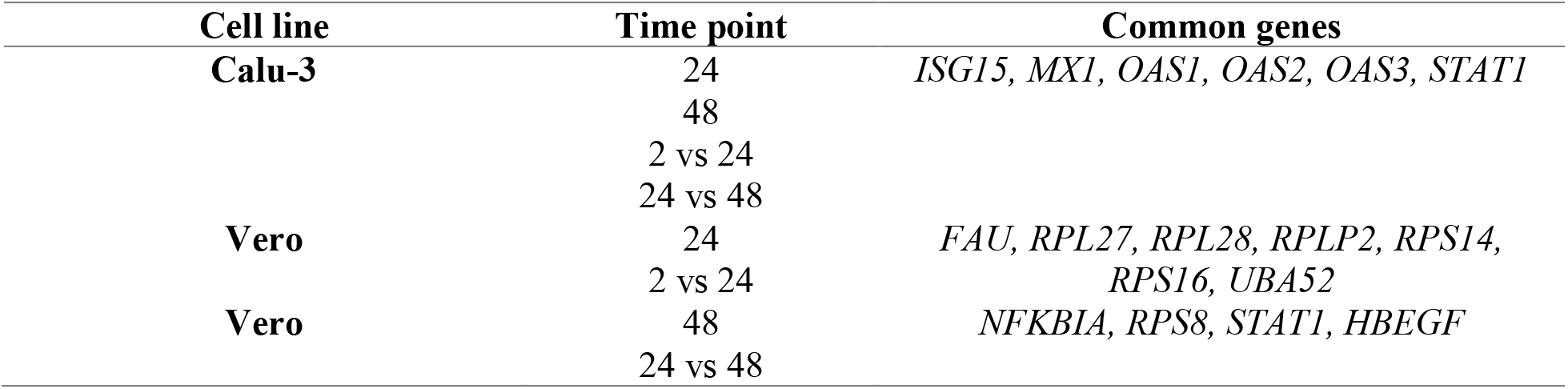
Genes involved in the *coronavirus pathway* shown via KEGG pathway analysis. Related to **Figure S2**.

### Lengths of host mRNA poly(A) tails change during SARS-CoV-2 infection

Polyadenylation has been previously shown in literature to be critical for many different cellular functions. The process promotes stabilisation of the RNA transcript (Beckel-Mitchener, 2002), trafficking into the cytoplasm (Fuke & Ohno, 2007), and translation into proteins (Park, Yi, Kim, Chang, & V, 2016). Furthermore, 3’ UTRs can include binding sites for RNA-binding proteins (RBPs) (Gebauer, Preiss, & Hentze, 2012) and microRNAs (miRNAs) (R. C. Lee, Feinbaum, & Ambros, 1993), which contribute to gene expression. However, only a small number of studies exploring changes in host poly(A) lengths during infections have been carried out to this date (Y. J. Lee & Glaunsinger, 2009). Therefore, we were interested in whether infection of cells with SARS-CoV-2 would elicit changes in polyadenylation of transcripts compared with control cells. The median poly(A) tail lengths of mitochondrial and non-mitochondrial transcripts were compared between control and infected cells at 2, 24 and 48 hpi with two different methods: *nanopolish* and *tailfindr*.

Firstly, *nanopolish* was used to analyse non-replicate direct RNA datasets (**Table 4**). Although the medians for each condition were similar in some datasets, we observed a significant (p < 0.05, Wilcoxon’s test of ranks two-tailed approach) poly(A) tail length increase in host non-mitochondrial RNA of infected cells compared with control cells in the 24 and 48 hpi in all susceptible cell lines. No significant change was observed in mitochondrial RNA.

**Table 4.** Significance of differential polyadenylation in Caco-2, Calu-3 and Vero cell lines measured by Wilcoxon’s test of ranks following *nanopolish* analysis of direct RNA datasets. Significant differential polyadenylation (compared with matched uninfected sample) was observed in all non-mitochondrial datasets (p < 0.05), except for the Calu-3 2 hpi dataset (in bold). Absence of significant differential polyadenylation in mitochondrial transcripts was also observed. Related to **Figure S3-4 & Tables S1-2**.

| Cell line | Time point | Type (Mitochondrial/Non-Mitochondrial) | Wilcoxon's test of ranks p-value |
| --- | --- | --- | --- |
| Caco-2 | 2 | Mitochondrial | W = 623, p-value = 0.5875 |
| Caco-2 | 24 | Mitochondrial | W = 647, p-value = 0.8504 |
| Caco-2 | 48 | Mitochondrial | W = 657, p-value = 0.4629 |
| **Caco-2** | **2** | **Non-mitochondrial** | **W = 89144178, p-value = 0.005541** |
| **Caco-2** | **24** | **Non-mitochondrial** | **W = 99425841, p-value = 0.00266** |
| **Caco-2** | **48** | **Non-mitochondrial** | **W = 91874180, p-value < 2.2e-16** |
| Calu-3 | 2 | Mitochondrial | W = 648, p-value = 1 |
| Calu-3 | 24 | Mitochondrial | W = 603, p-value = 0.9163 |
| Calu-3 | 48 | Mitochondrial | W = 625, p-value = 0.4283 |
| Calu-3 | 2 | Non-mitochondrial | W = 86343421, p-value = 0.06655 |
| **Calu-3** | **24** | **Non-mitochondrial** | **W = 92324958, p-value < 2.2e-16** |
| **Calu-3** | **48** | **Non-mitochondrial** | **W = 56602100, p-value < 2.2e-16** |
| Vero | 2 | Mitochondrial | W = 647, p-value = 0.5388 |
| Vero | 24 | Mitochondrial | W = 686, p-value = 0.3935 |
| Vero | 48 | Mitochondrial | W = 565, p-value = 0.7233 |
| **Vero** | **2** | **Non-mitochondrial** | **W = 36539892, p-value = 1.149e-05** |
| **Vero** | **24** | **Non-mitochondrial** | **W = 32813975, p-value = 3.194e-15** |
| **Vero** | **48** | **Non-mitochondrial** | **W = 34564286, p-value = 1.505e-09** |

As *nanopolish* results only used data from non-replicate direct RNA datasets, a second approach was utilised. This involved direct cDNA datasets with triplicates for each condition using *tailfindr* to confirm the results of *nanopolish* at the gene level. As the direct cDNA dataset is double-stranded, either strand of the cDNA can be sequenced. Therefore, information on both poly(A) and poly(T) lengths were obtained, which were weakly correlated (**Figure S3 & Table S1**). When the number of differentially polyadenylated transcripts between control and infected cells were compared with *nanopolish* and *tailfindr* poly(A) and poly(T) methods with the Calu-3 48 hpi non-mitochondrial data (Wilcoxon’s test, padj < 0.05), the *tailfindr* poly(A) dataset showed no significant differential polyadenylation (**Table S2**). The lack of significance in the *tailfindr* poly(A) data can be explained by the fact that less data was available from the poly(A) dataset compared to the poly(T) dataset. The full-length reads were comprised of 0.5% of poly(A) and 99.5% of poly(T) strands, which made up ~58% of the total number of detected reads with valid Ensembl ID’s. This may be attributed to the process of ONT direct cDNA sequencing, where the motor protein is situated on the 5’ end of each strand. This means that the poly(T) sequence is sequenced first, whereas the poly(A) sequence is sequenced last for each respective strand. Therefore, this method of sequencing would lead to higher quantity and accuracy of poly(T) sequences compared with poly(A) sequences (**Table S2**). To test this proposition, we compared the median lengths of poly(A/T) tails per gene between *nanopolish* and *tailfindr* Calu-3 48 hpi datasets via the Spearman’s correlation test. Weak significant positive correlations between *nanopolish* poly(A) and *tailfindr* poly(T) datasets were observed (r = 0.12-0.27, p-value < 0.05). In contrast, *nanopolish* poly(A) and *tailfindr* poly(A) data were not significantly correlated (**Figure S4**), leading us to choose poly(T) length as a proxy for direct RNA-inferred poly(A) length.

### Transcripts of ribosomal protein genes are elongated during SARS-CoV-2 infection

Specific genes involved in differential polyadenylation were investigated using a linear mixed-model method using *tailfindr* outputs. The most interesting dataset was Calu-3 48 hpi, where twelve genes were found to be significantly increased in poly(A) length (up to ~101 nt in mean poly(A) length) in the infected cells compared with control cells (*UQCRC1*, *RPL30*, *RPS12*, *RPL13*, *KRT17*, *CXCL8*, *RPS6*, *ZBTB44*, *MIEN1*, *RPS4X*, *RPL10 and* a lncRNA-*ENSG00000273149*) (**Figure S5**). Using *multiGO*, GO biological terms of genes with increased poly(A) length were found, which included *viral transcription* (*RPS12*, *RPL30*, *RPS6*, *RPL13*, *RPS4X*, *RPL10*) (enrichment p-value < 0.0001) (**Figure 4 & Data S3**). KEGG pathways of these genes included the *coronavirus disease* and *ribosome* pathways (**Figure S6 & Data S3**). This suggests that poly(A) tail elongation may be directly linked to SARS-CoV-2 infections, as opposed to a randomly occurring event. A small number of mitochondrial genes were also found to be differentially polyadenylated (including *ENSG00000198888/MT-MD1*) in both Calu-3 and Vero cell lines. Additionally, among the twelve genes which were found to be increased in poly(A) length in *tailfindr* mixed-model analysis, eight genes were also found to be significantly increased in poly(A) length in *nanopolish* analysis after log-transformation and p-value adjustment (*ENSG00000273149*, *RPS12*, *RPL30*, *RPS6*, *RPL13*, *MIEN1*, *RPS4X*, *RPL10*, padj < 0.05). This confirmed the robustness of these results, which increased the confidence of true poly(A) elongation in these eight genes. The other four genes were unable to be detected in *nanopolish* datasets even when padj thresholds were relaxed.

**Figure 4.**
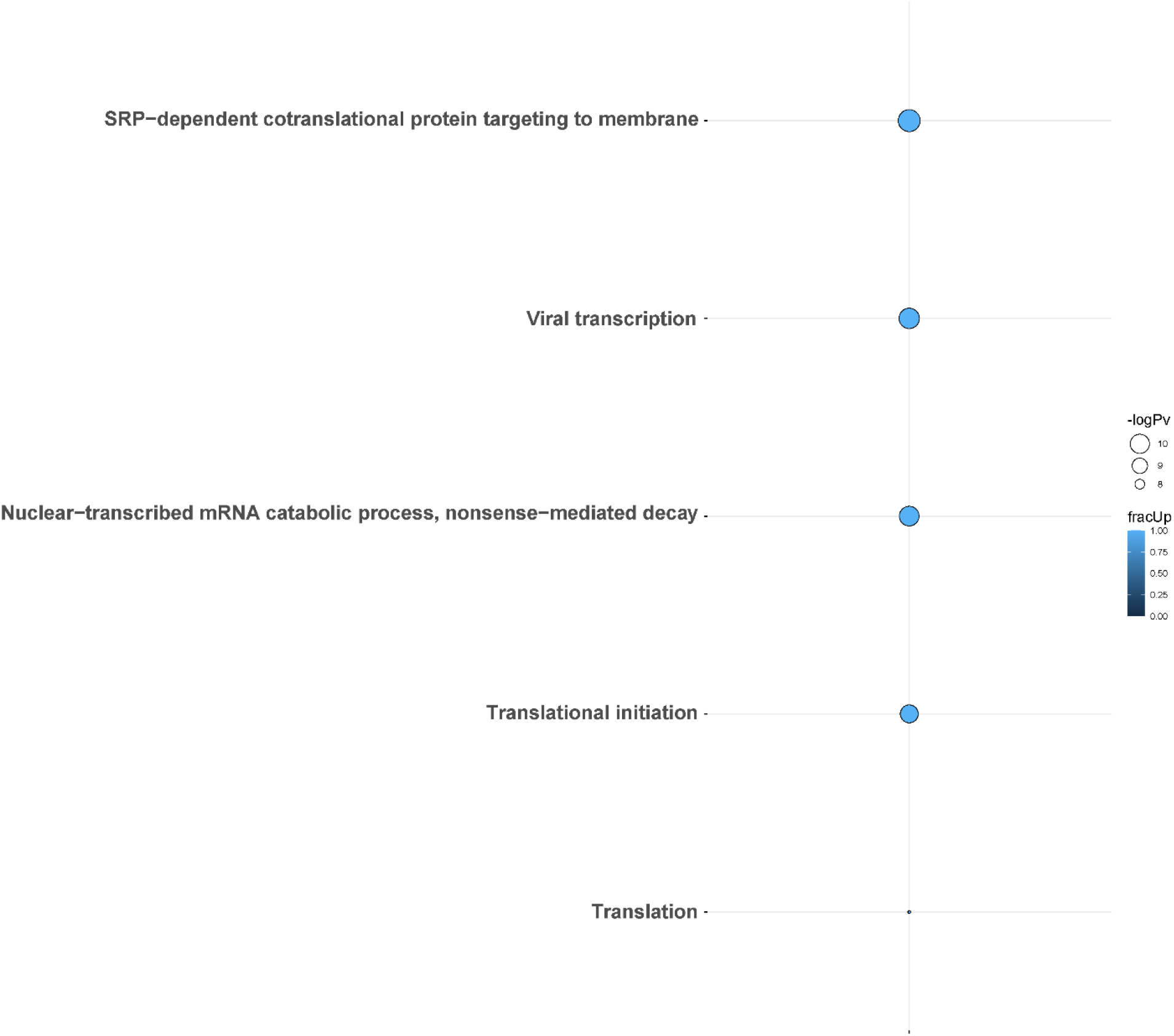
GO biological terms from genes with differential poly(A) tail length in *tailfindr* poly(T) mixed-model analyses from the Calu-3 48 hpi direct cDNA dataset. Genes involved in *viral transcription*, *translation*, *translational initiation*, *SRP-mediated cotranslational protein targeting to membrane and nuclear-transcribed mRNA catabolic process* and *nonsense-mediated decay* pathways were increased in poly(A) tail length after infection. The bubble size and colour indicate the −log10 enrichment p-values and the fraction of genes with increased polyadenylation, respectively. Thresholds of padj < 0.05, enrichment p-value < 0.0001 were used. Only bubbles which meet the thresholds are shown. Related to **Figures S5-6 & Data S3**.

### Ribosomal protein genes *RPS4X* and *RPS6* show increased poly(A) tail lengths and downregulated expression upon infection

We next investigated whether there was any relationship between differential polyadenylation and differential expression results in response to infection. Interestingly, when comparing the GO terms which were shared among the differential polyadenylation and differential expression results of Calu-3 48 hpi direct-cDNA datasets, the results showed an apparent correlation between the two analyses. The enriched GO terms were composed of genes with mainly increased poly(A) tail lengths and decreased expression levels after infection (**Figure 5**). Upon closer inspection, we found that many of the genes involved in these GO terms were associated with the ribosome. Of note, two genes (*RPS4X* and *RPS6*) which contributed to all GO terms, both showed an increase in poly(A) length and downregulated gene expression. This overlap was significant (hypergeometric test, p=0.018). When the KEGG pathways were compared in a similar manner, we observed that the *coronavirus disease* pathway was shared between differential polyadenylation and expression datasets (**Figure S7**). Unlike the GO terms, most genes had an increase in poly(A) tail length but were upregulated in differential expression levels, although many ribosomal genes were downregulated in the same dataset. For example, *CXCL8* had an increased poly(A) length after infection and was upregulated in the expression level results, unlike the ribosomal protein genes described above. This suggested that the correlation between increased poly(A) tails and decreased expression levels were shown specifically in the ribosome-related protein genes and indicated the importance of ribosomal protein genes during SARS-CoV-2 infections.

**Figure 5.**
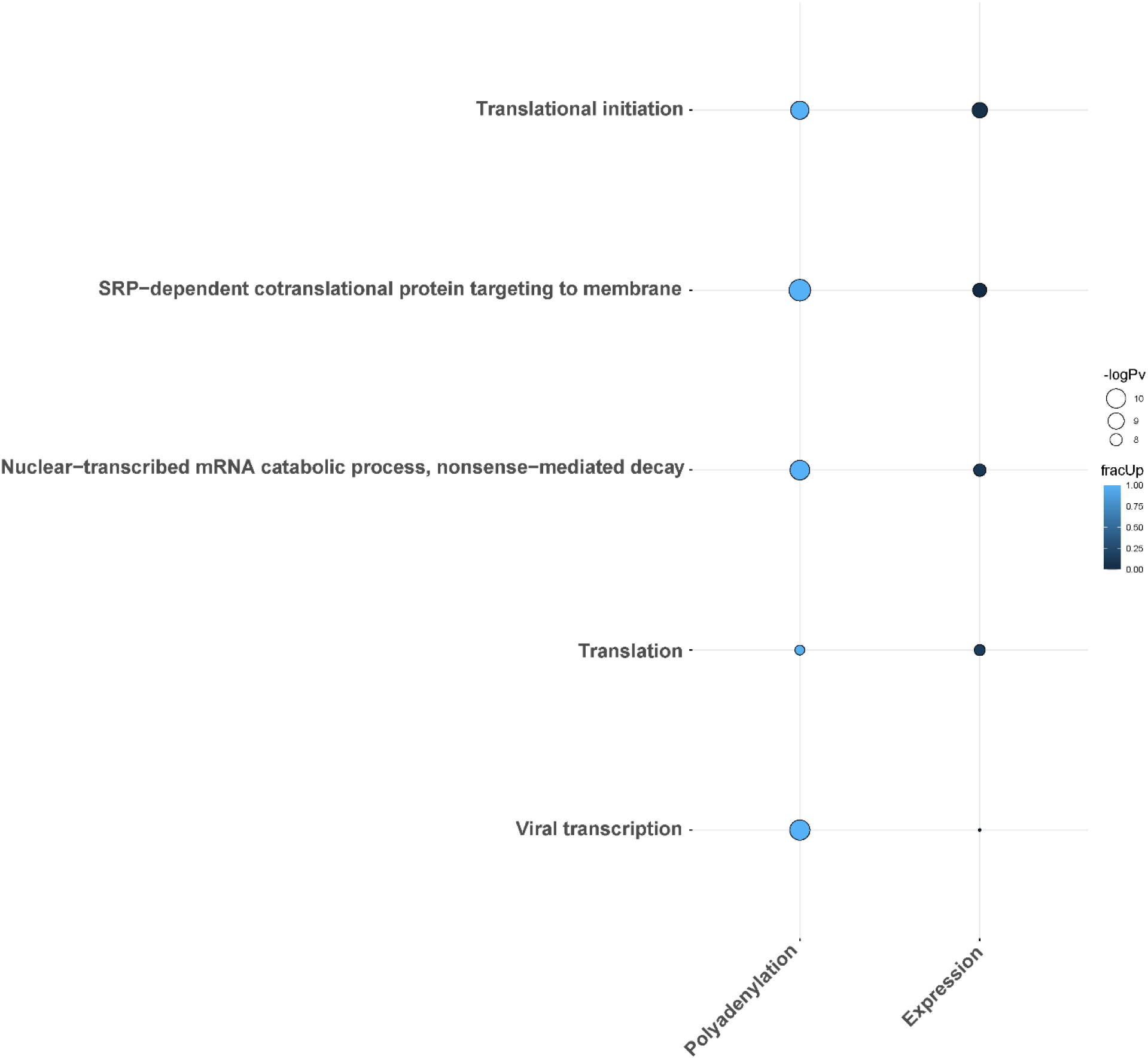
Direction of differentially polyadenylated and expressed genes belonging to common GO biological terms in the two analysis methods using the Calu-3 48 hpi direct cDNA dataset. The plot shows a potential correlation in increased poly(A) tail length and downregulation in gene expression. The bubble size and colour indicate the −log10 enrichment p-values and fraction of upregulated genes/genes with increased polyadenylation, respectively. Thresholds of padj < 0.05, enrichment p-value < 0.0001 were used. Only bubbles which meet the thresholds are shown. Related to **Figures S7 & Data S4**.

### Differential transcript usage occurs between control and infected cells during SARS-CoV-2 infection

Differential transcript usage is the differential presence of transcripts between different conditions measured via identifying the proportion of each transcript against the total pool of transcripts and is another valuable feature of ONT RNA-seq. Using *DRIMSeq* and *StageR*, significant differential transcript usage was observed in all three SARS-CoV-2 susceptible cell lines (Calu-3, Caco-2 and Vero) between infected and mock-control cells. These events were observed in three time points (2, 24, 48 hpi) in Caco-2, two time points in Calu-3 (2, 48 hpi) and one time point in Vero cells (24 hpi). This included a processed transcript - *SLC37A4-205* - in the Caco-2 2 hpi dataset and a retained intron transcript - *GSDMB-208* – in the Calu-3 48 hpi dataset (**Figure 6 & Table 5**). These results suggested that non-protein-coding transcripts could also show differential usage as with protein-coding transcripts. Furthermore, these results revealed that differential transcript usage events were not specific to a given time point and may have also occurred in a cell-specific manner.

**Figure 6.**
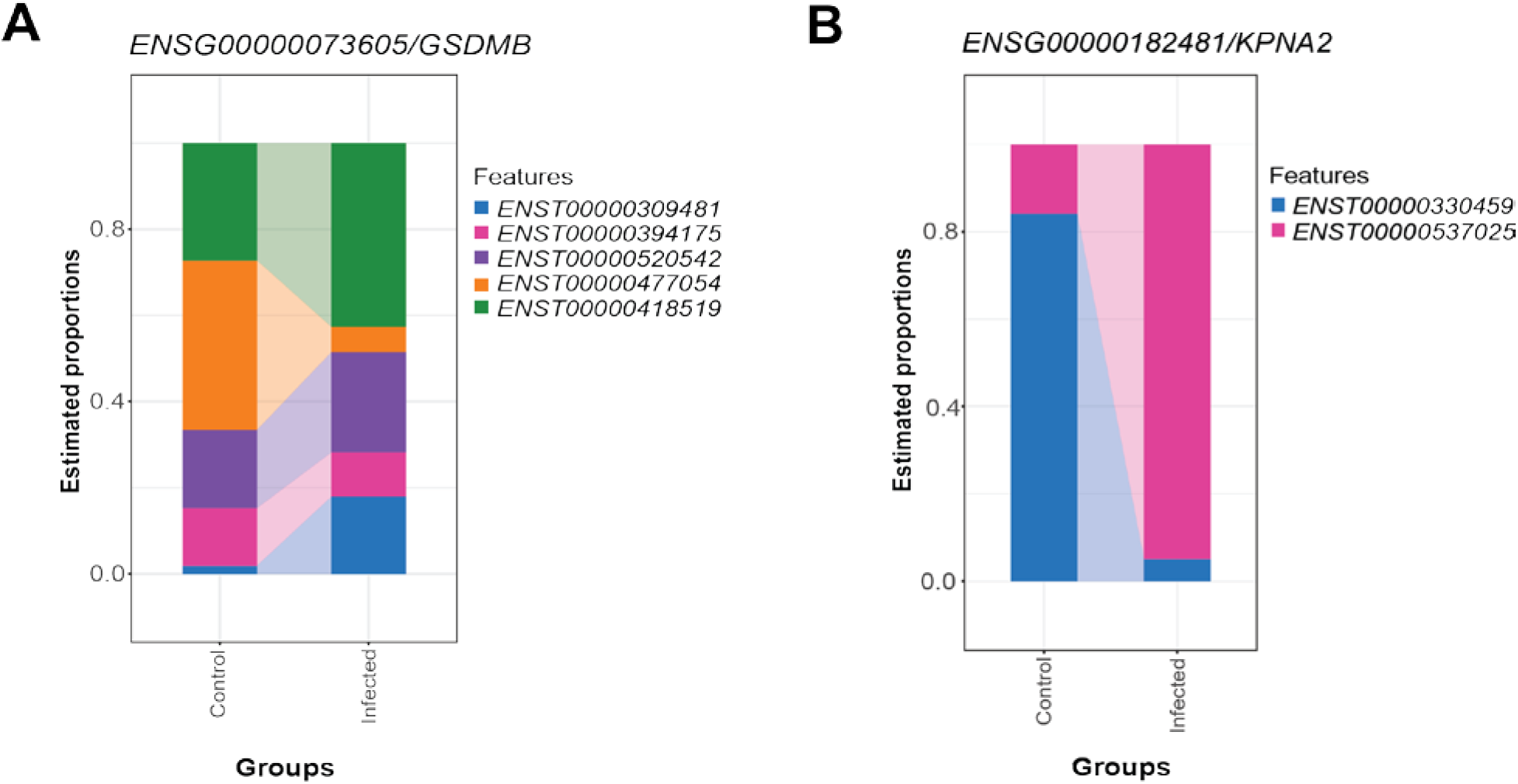
Differential transcript usage in the Calu-3 48 hpi dataset. **A)** Differential estimated proportions of transcripts of *ENSG00000073605/GSDMB* between control and infected cells. **B)** Differential estimated proportions of transcripts of *ENSG00000182481/KPNA2* between control and infected cells. Related to **Table 5**.

**Table 5.** List of isoforms with significant differential usage in Caco-2, Calu-3 and Vero. Related to **Figure 6**.

| Cell line | Time point | Isoforms with differential transcript usage |
| --- | --- | --- |
| Caco-2 | 2 | *IPO5-201, IPO5-230, SLC37A4-205, SLC37A4-217* |
| Caco-2 | 24 | *NACA-221, SERF2-201* |
| Caco-2 | 48 | *PKM-204, RPL4-201, RPL4-215* |
| Calu-3 | 2 | *ATP13A3-201, STIL-201, STIL-202, TRA2A-202, TRA2A-211* |
| Calu-3 | 48 | *SPTBN1-207, GSDMB-208, KPNA2-201, KPNA2-202, RAE1-202, RAE1-203, ATIC-205, ATIC-207, ADK-202, ADK-206, ATRAID-201, ATRAID-207, ANKRD12-201, ANKRD12-203, WARS1-202, WARS1-204* |
| Vero | 24 | *ARPC1B-202, ARPC1B-201* |

## Discussion

Long-read sequencing enabled the detection of differential polyadenylation, transcript usage and gene expression level changes within *in vitro* SARS-CoV-2 infection models. Firstly, median poly(A) tail lengths between control and infected cells in direct RNA data were estimated using *nanopolish*. This showed that the non-mitochondrial median poly(A) tail lengths were significantly increased in all three cell lines (Caco-2, Calu-3 and Vero) at the 24 and 48 hpi (**Table 4**). These results suggested that infection with SARS-CoV-2 may cause an increase in the poly(A) lengths of non-mitochondrial transcripts. We explored this further using *tailfindr*. The results from the mixed-effects model analysis showed poly(A) tail elongation after infection in Calu-3 and Vero cells at the 48 hpi. In Calu-3 cells, six genes were involved in viral transcription (*RPS12*, *RPL30*, *RPS6*, *RPL13*, *RPS4X*, *RPL10*) (**Figure 6**). This indicated that polyadenylation may play a role in aiding the virus to generate viral mRNA for further protein production or to replicate during infection with SARS-CoV-2. This group of genes is involved in the formation of the ribosome, which is required for protein synthesis. The result is relevant as the SARS-CoV-2 non-structural protein Nsp1 binds to the 40S subunit of the ribosome and inhibits translation initiation of cellular mRNA (Lapointe et al., 2021; Schubert et al., 2020). Ribosomal proteins have also been known to be associated with viral transcription/replication as the host ribosomal machinery needs to be utilised to produce viral proteins for these processes (Stern-Ginossar, Thompson, Mathews, & Mohr, 2019). Therefore, further investigation is warranted to explore the link between increased polyadenylation in host cells after infection with SARS-CoV-2. It would also be of interest to study whether host defense ability decreases with elevated polyadenylation of transcripts related to viral transcription and the ribosome. A lncRNA and a small number of mitochondrial transcripts were also observed with an elongated poly(A) length in infected cells compared with control cells. This is of interest as it suggests that not only the protein-coding genes may be able to play a role in host responses to SARS-CoV-2. Furthermore, as elongated poly(A) tails were observed at a late time point (48 hpi) in Vero cells, it suggested that differential polyadenylation may be more likely to occur at later stages of infection compared with early stages.

We also explored the observation that many of the genes involved in the commonly enriched GO terms and KEGG pathways were increased in poly(A) tail length and decreased in gene expression (**Figures 5 & S7**). Interestingly, the majority of genes which were involved in both of these observations were ribosomal proteins, such as *RPS4X* and *RPS6* which encode for proteins in the 40S ribosomal subunit. In contrast, a non-ribosomal gene *CXCL8* had increases in both poly(A) length and expression level after infection, which suggested that this correlation belonged exclusively to the ribosomal protein genes. This was an interesting observation as decreased expression of ribosomal proteins in response to SARS-CoV-2 infections have been observed previously (Lieberman et al., 2020), due to the effect of global suppression of ribosomal activity initiated by the virus. However, the increase in poly(A) lengths in the same transcripts was unexpected, as elongation of poly(A) tails are indicative of increase in stability, in contrast to the decrease in expression levels. Why these observations were uniquely presented in these ribosomal protein genes is currently unclear. However, we speculate this may be due to the competition between viral-driven expression downregulation and host-driven post-transcriptional regulation for increased stability of mRNA. In some cases, aberrant polyadenylation has been linked to aid the destruction of eukaryotic mRNA (Y. J. Lee & Glaunsinger, 2009), which may provide an alternative explanation for this phenomenon.

Supporting the importance of the ribosome during SARS-CoV-2 infection, the *translation* GO term was enriched in infected Calu-3 and Vero cells (**Figure 3**). Among the genes involved in translation, the *EIF1* gene encodes for the eukaryotic translation initiation factor 1, which partakes in translation initiation in eukaryotes by forming a part of the 43S preinitiation complex along with the 40S ribosomal subunit. As mentioned earlier, according to Lapointe et al., (2021), this factor may enhance the binding of SARS-CoV-2 Nsp1 protein to the host 40S subunit, perhaps via changing the conformation of the mRNA entry channel. This facilitates host translation inhibition by competing with host mRNA, which has been shown to also decrease translation of viral mRNA. However, another study showed that the binding of Nsp1 to the 40S subunit induced preferential translation of viral mRNA over host mRNA (Schubert et al., 2020). These findings may explain the downregulation the *EIF1* as a host response to viral infection.

Differential transcript usage has not yet been extensively studied with SARS-CoV-2 infections but has been useful for studying other illnesses like Parkinson’s disease (Dick et al., 2020). Differential transcript usage was observed in all three SARS-CoV-2 susceptible cell lines studied – Caco-2, Calu-3 and Vero, where protein-coding, processed and retained-intron transcripts were involved. Calu-3 48 hpi data showed the greatest number of genes that had undergone differential transcript usage, where a retained-intron transcript *GSDMB-208* showed differential usage in infected cells compared with control cells (**Figure 6**). SARS-CoV-2 induces pyroptosis in human monocytes (Ferreira et al., 2021), and the Gasdermin family of proteins has been implicated in cell death where the granzyme-mediated cleavage of GSDMB can activate pyroptosis (Z. Zhou et al., 2020). Furthermore, *KPNA2* transcripts also showed differential usage in Calu-3 cells at 48 hpi. *KPNA2* is an importin that is bound by ORF6 of the virus to block nuclear IRF3 and ISGF3 to antagonise IFN-1 production and signalling (Xia et al., 2020). This suggested that transcripts with differential usage may be involved in important pathways contributing to host responses towards viral infection or the evasion of these responses by the virus. Hence, the specific activity of each transcript as opposed to the activity at the gene level should be further investigated.

Overall, our long-read sequencing datasets agreed with differential expression studies in the literature. In agreement with previous studies, our differential expression analysis using direct cDNA datasets showed varied host gene expression activity upon infection in different cell types (**Figure 1**). Although Vero cells are imperfect *in vitro* models for SARS-CoV-2 infection, we note that the *ROS metabolic pathway* was enriched in our direct cDNA data across the SARS-CoV-2 susceptible cell lines (**Figures 3 & S2**). The mitochondria can be linked to the ROS metabolic pathway as it produces ROS which can induce increased oxidative stress in cells, potentially leading to cell death (Orrenius, 2007). These results increased support for the idea that mitochondrial processes are important during these infections (Codo et al., 2020; Singh et al., 2021).

We also observed the downregulation of pathways involved in neurological pathologies such as *Parkinson’s disease* (Martinelli et al., 2021) (**Figure S2**). The fact that some enriched pathways were involved in non-respiratory, neurological illnesses suggests potential modes of action for SARS-CoV-2 co-morbidities. There are now increasing numbers of studies reporting on the relevance between COVID-19 and other diseases. This includes clinical data where patients with COVID-19 can develop neurological problems which are not only non-specific (e.g. headaches), but also varied, including such maladies as viral meningitis, encephalitis (Moriguchi et al., 2020), olfactory and gustatory dysfunction (Luers et al., 2020), and dementia-related symptoms similar to Alzheimer’s disease (Y. Zhou et al., 2021). However, the relevance of this pathway in non-neuronal cells is potentially limited.

## Limitations & future directions

In our study, we utilised *in vitro* models of SARS-CoV-2 infections using continuous cell lines with a low MOI of 0.1, which may hinder the biological relevance of these results. However, our results confirmed that the use of ONT RNA-seq methods enabled the detection of full-length isoforms, differential polyadenylation and transcript usage. This provides evidence to pursue further investigations with more sophisticated models such as air-liquid-interface cultured organoids from healthy human nasal swabs or *in vivo* models such as ferrets. The ideal MOI should also be found via optimisation studies.

Moreover, our results showed that the *nanopolish* and *tailfindr* methods had significant weak positive correlations (r < 0.3, p-value < 0.05) in the median poly(A) lengths from *nanopolish* and median poly(T) lengths from *tailfindr* (**Figure S4**). However, the median poly(A) lengths of *tailfindr* showed non-significant correlations with *nanopolish* poly(A) data. The increased significance of poly(T) transcripts may have occurred because more data was available from the poly(T) dataset. As our results supported similar findings from Krause et al. (2019), we speculate that these discrepancies between poly(A) and poly(T) datasets using ONT direct cDNA sequencing may arise in future studies. We acknowledge that our data is preliminary and the correlation between *nanopolish* and *tailfindr* data should be tested via direct RNA datasets with replicates to validate these findings.

Functional work should also be carried out to further validate the results of this study. For differential expression analysis, knock-down experiments within the same cell lines using CRISPR technology may be utilised to evaluate the effects of differentially expressed genes identified in this study. Furthermore, functional work for polyadenylation may be approached by utilising cell lines with plasmids containing gene sequences of interest followed by a poly(A) sequence of varying lengths. Assays such as measuring viral titre and further RNA-seq may be used to test the effects of these alterations after infection with SARS-CoV-2.

## Conclusions

Overall, by utilising three ONT RNA-seq methodologies, we generated an in-depth characterisation of differential expression, polyadenylation and differential transcript usage of cell lines infected *in vitro* by SARS-CoV-2. Unravelling the pathways associated with duration of infection and responses of differing cell types using long-read methods will provide novel insights into the pathogenesis of SARS-CoV-2.

## Funding

This research was supported by a University of Melbourne “Driving research momentum” award (to LC) and NHMRC EU project grant (GNT1195743 to LC). KS was supported by an NHMRC Investigator grant. The Melbourne WHO Collaborating Centre for Reference and Research on Influenza was supported by the Australian Government Department of Health. MC was supported by an Australian National Health and Medical Research Council Investigator Fellowship (APP1196841). JC was supported by the Miller Foundation and the Australian Government Research Training Programme (RTP) scholarship.

## Supplementary items

**Data S1. Relative percentages of mapped reads to host, virus or sequin genomes in all datasets**

**Data S2. *multiGO* outputs of GO biological and KEGG analyses from the direct cDNA differential expression results, including −log10 p-values from the enrichment analyses and genes involved in the GO terms/KEGG pathways**

**Data S3. *multiGO* outputs of GO biological and KEGG analyses from the Calu-3 48 hpi direct cDNA differential polyadenylation results, including −log10 p-values from the enrichment analyses and genes involved in the GO terms/KEGG pathways**

**Data S4. *multiGO* outputs of GO biological and KEGG analyses from the Calu-3 48 hpi direct cDNA differential polyadenylation vs expression results, including −log10 p-values from the enrichment analyses and genes involved in the GO terms/KEGG pathways**

**Table S1. Correlations between *tailfindr* median poly(A) and poly(T) lengths per gene using direct cDNA data.** Weak positive correlations were observed between poly(A) and poly(T) datasets (r < 0.4), with all correlations being significant (p-value < 0.05, Pearson’s correlation test). Related to **Figure S3 & Table 4**.

**Table S2. Comparison of significantly differentially polyadenylated non-mitochondrial gene clusters in the Calu-3 48 hpi datasets between *nanopolish, tailfindr* poly(A) and poly(T) outputs (before log-transformation) using Wilcoxon’s test of ranks.** *nanopolish* and *tailfindr* poly(T) results were more comparable compared with *tailfindr* poly(A) results, as no significant polyadenylation was observed in *tailfindr* poly(A) data. Related to **Figure S3-4 & Table 4**.

**Figure Sl. Volcano plots showing changes in differential expression between control and infected cells across two time points in direct cDNA datasets using *DESeq2* and the interactive term.** X-axis represents log2FC and Y-axis displays −log10 padj. 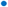 padj < 0.05, 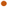 |log2(FC)| > 1, 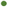 both. Plots reveal increase in differential expression at latter time points compared with earlier time points in Caco-2, Calu-3 and Vero cell lines, and a lack of significant differential expression was observed between A549 0 vs 24 hpi. Related to **Figure 1**.

**Figure S2. KEGG pathways from differentially expressed genes in Calu-3, Caco-2 and Vero direct cDNA datasets analysed by *DESeq2***. Results include datasets (in order); Caco-2 24 vs 48, Caco-2 48, Calu-3 2 vs 24, Calu-3 24 vs 48, Calu-3 24, Calu-3 48, Vero 2 vs 24, Vero 24 vs 48, Vero 24 and Vero 48 hpi. Strongest KEGG pathway enrichment was observed in the Calu-3 cells including pathways such as *influenza A*, *measles* and *coronavirus diseas*e. The bubble size and colour indicate the −log10 enrichment p-values and fraction of upregulated genes, respectively. Thresholds of padj < 0.05 and enrichment p-value < 0.0001 in at least one dataset were used for generating the plot, and all terms with padj < 0.05 and enrichment p-value < 0.05 were deemed as significant for the analysis. Related to **Figure 3 & Data S2**.

**Figure S3. Correlations between median poly(A/T) lengths from *tailfindr* analyses.** Scatter plots comparing *tailfindr* poly(A) vs poly(T) datasets from Caco-2 **A)** 2 hpi, **B)** 24 hpi, **C)** 48 hpi, Calu-3 **D)** 2 hpi, **E)** 24 hpi, **F)** 48 hpi, Vero **G)** 2 hpi, **H)** 24 hpi**, I)** 48 hpi. Weak correlation between median poly(A) and poly(T) lengths were observed (R < 0.4, exact values along with p-values are indicated in **Table S1**), where each dot represents a gene. Related to **Tables 4 & S1-2**.

**Figure S4. Correlations between median poly(A) lengths from *nanopolish* and poly(A/T) lengths from *tailfindr* analyses.**

Scatter plots comparing **A)** median *nanopolish* poly(A) and *tailfindr* poly(A) lengths in control cells **B)** median *nanopolish* poly(A) and *tailfindr* poly(T) lengths in control cells, **C)** median *nanopolish* poly(A) and *tailfindr* poly(A) lengths in infected cells and **D)** median *nanopolish* poly(A) and *tailfindr* poly(T) lengths in infected cells. Weak significant positive correlations were observed between the *nanopolish* poly(A) and *tailfindr* poly(T) datasets (r = 0.12-0.27, p-value < 0.05, Spearman’s correlation test). No significant correlation was observed between *nanopolish* poly(A) and *tailfindr* poly(A) datasets (p-value > 0.05, Spearman’s correlation test). Related to **Tables 4 & S2**.

**Figure S5. Raincloud plots of raw, untransformed poly(T) tail lengths in twelve genes with increased poly(A) tail length in the *tailfindr* poly(T) mixed-model analysis with log-transformation.** Genes include **A)** *UQCRC1*, **B)** *RPL30*, **C)** *RPS12*, **D)** *RPL13*, **E)** *KRT17*, **F)** *CXCL8*, **G)** *RPS6*, **H)** *ZBTB44*, **I)** *MIEN1*, **J)** *RPS4X*, **K)** *RPL10* and **L)** a lncRNA. Related to **Figure 4**.

**Figure S6. KEGG pathways from genes with differential poly(A) tail length in *tailfindr* poly(T) mixed-model analyses from the Calu-3 48 hpi dataset.** Genes involved in *coronavirus disease* and *ribosome* pathways were increased in poly(A) length after infection. The bubble size and colour indicate the −log10 enrichment p-values and the fraction of genes with increased polyadenylation, respectively. Thresholds of padj < 0.05, enrichment p-value < 0.0001 were used. Only bubbles which meet the thresholds are shown. Related to **Figure 4 & Data S3**.

**Figure S7. KEGG pathways of differentially polyadenylated and expressed genes from the Calu-3 48 hpi direct cDNA dataset.** Only the *coronavirus disease* pathway was shared between the two analyses. The plot shows a potential correlation in increased poly(A) tail length and downregulation in gene expression. The bubble size and colour indicate the −log10 enrichment p-values and fraction of upregulated genes/genes with increased polyadenylation, respectively. Thresholds of padj < 0.05, enrichment p-value < 0.0001 were used. Related to **Figure 5 & Data S4**.

## Notes

https://www.ncbi.nlm.nih.gov/bioproject/PRJNA675370

### Competing Interest Statement

The authors have declared no competing interest.

