## Supplementary Tables and Figures for "Long-read RNA sequencing identifies polyadenylation elongation and differential transcript usage of host transcripts during SARS-CoV-2 *in vitro* infection"

**Table S1. Correlations between *tailfindr* median poly(A) and poly(T) lengths per gene using direct cDNA data.** Weak positive correlations were observed between poly(A) and poly(T) datasets (r < 0.4), with all correlations being significant (p-value < 0.05, Pearson's correlation test). Related to **Figure S3 & Table 4**.

| Cell line | Time point | r | P-value |
|---|---|---|---|
| **Caco-2** | 2 | 0.247 | $< 2.2 \times 10^{-16}$ |
| **Caco-2** | 24 | 0.208 | $1.873 \times 10^{-7}$ |
| **Caco-2** | 48 | 0.302 | $< 2.2 \times 10^{-16}$ |
| **Calu-3** | 2 | 0.233 | $< 2.2 \times 10^{-16}$ |
| **Calu-3** | 24 | 0.213 | $1.785 \times 10^{-13}$ |
| **Calu-3** | 48 | 0.267 | $<2.2 \times 10^{-16}$ |
| **Vero** | 2 | 0.370 | $< 2.2 \times 10^{-16}$ |
| **Vero** | 24 | 0.249 | $2.043 \times 10^{-11}$ |
| **Vero** | 48 | 0.279 | $2.524 \times 10^{-14}$ |

**Table S2. Comparison of significantly differentially polyadenylated non-mitochondrial gene clusters in the Calu-3 48 hpi datasets between *nanopolish*, *tailfindr* poly(A) and poly(T) outputs (before log-transformation) using Wilcoxon's test of ranks.** *nanopolish* and *tailfindr* poly(T) results were more comparable compared with *tailfindr* poly(A) results, as no significant polyadenylation was observed in *tailfindr* poly(A) data. Related to **Figure S3-4 & Table 4**.

| Direction of Polyadenylation | ***nanopolish*** (nonMT) | ***tailfindr* poly(A)** (nonMT) | ***tailfindr* poly(T)** (nonMT) |
|---|---|---|---|
| Up | 457 | 0 | 72 |
| Down | 3 | 0 | 1 |
| Total | 460 | 0 | 73 |

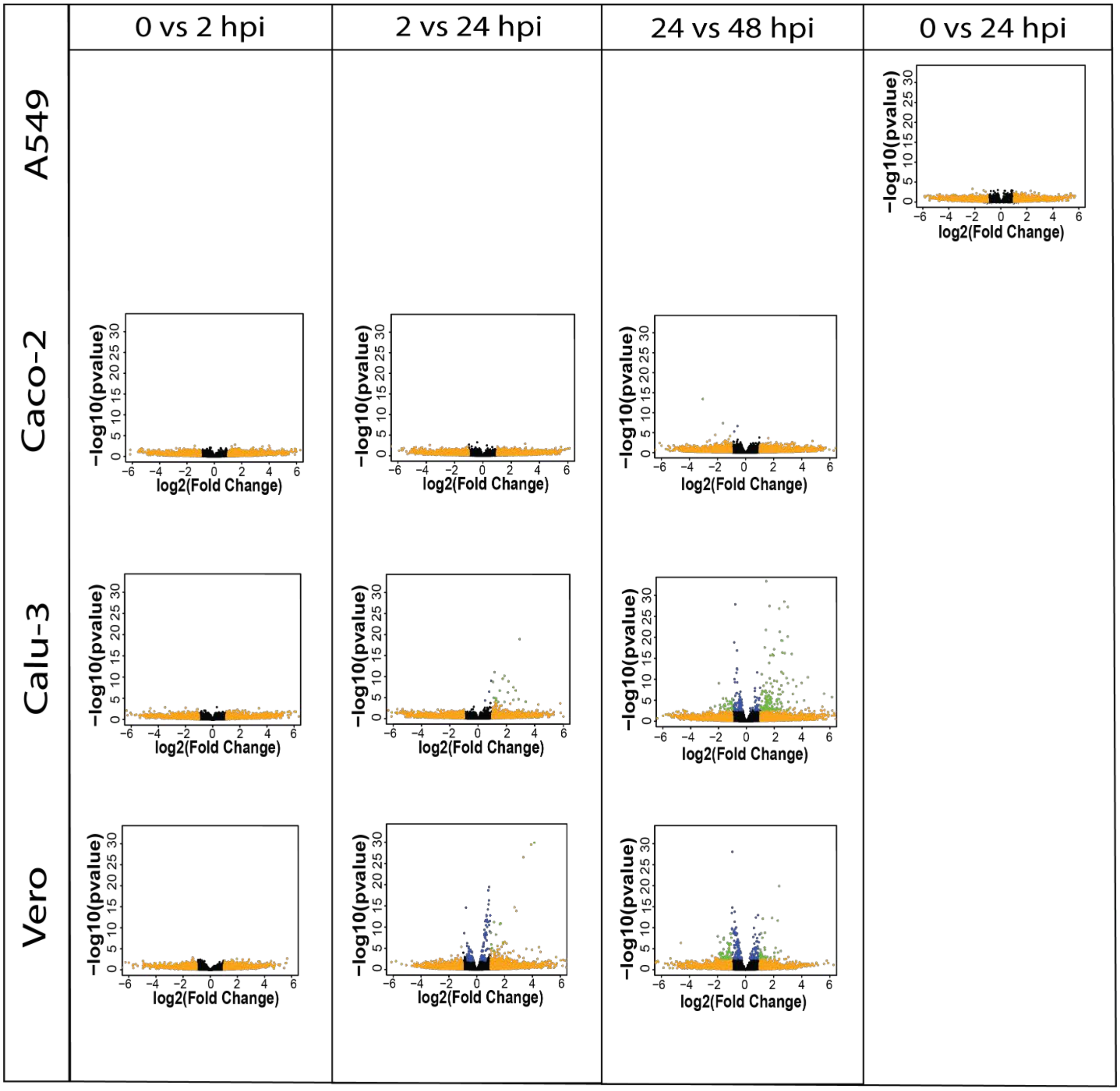


**Figure S1. Volcano plots showing changes in differential expression between control and infected cells across two time points in direct cDNA datasets using *DESeq2* and the interactive term.** X-axis represents log2FC and Y-axis displays –log10 padj. • padj < 0.05, • |log2(FC)| > 1, • both. Plots reveal increase in differential expression at latter time points compared with earlier time points in Caco-2, Calu-3 and Vero cell lines, and a lack of significant differential expression was observed between A549 0 vs 24 hpi. Related to **Figure 1**.

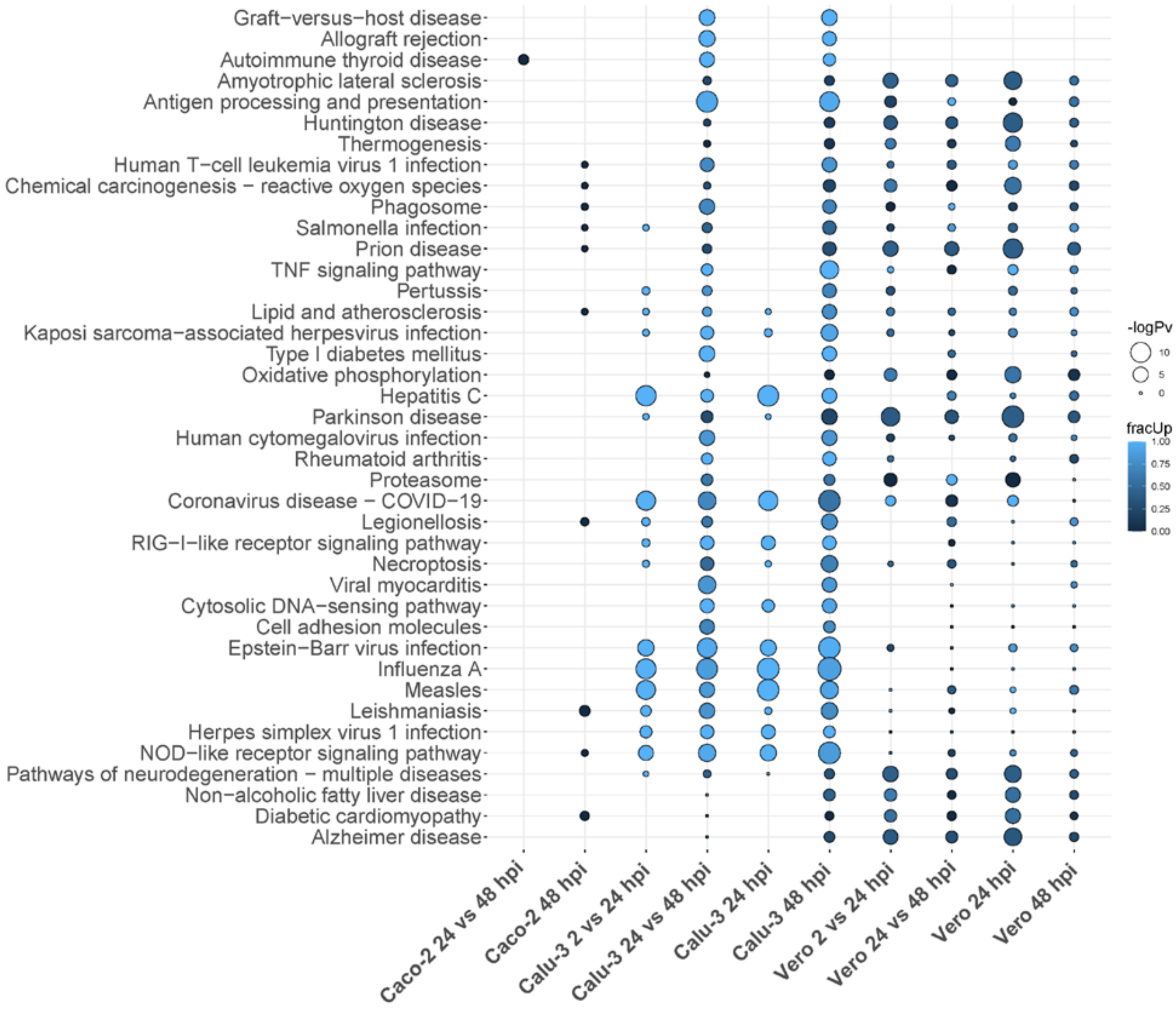


**Figure S2. KEGG pathways from differentially expressed genes in Calu-3, Caco-2 and Vero direct cDNA datasets analysed by *DESeq2***. Results include datasets (in order); Caco-2 24 vs 48, Caco-2 48, Calu-3 2 vs 24, Calu-3 24 vs 48, Calu-3 24, Calu-3 48, Vero 2 vs 24, Vero 24 vs 48, Vero 24 and Vero 48 hpi. Strongest KEGG pathway enrichment was observed in the Calu-3 cells including pathways such as *influenza A*, *measles* and *coronavirus disease*. The bubble size and colour indicate the -log10 enrichment p-values and fraction of upregulated genes, respectively. Thresholds of padj < 0.05 and enrichment p-value < 0.0001 in at least one dataset were used for generating the plot, and all terms with padj < 0.05 and enrichment p-value < 0.05 were deemed as significant for the analysis. Related to **Figure 3 & Data S2**.

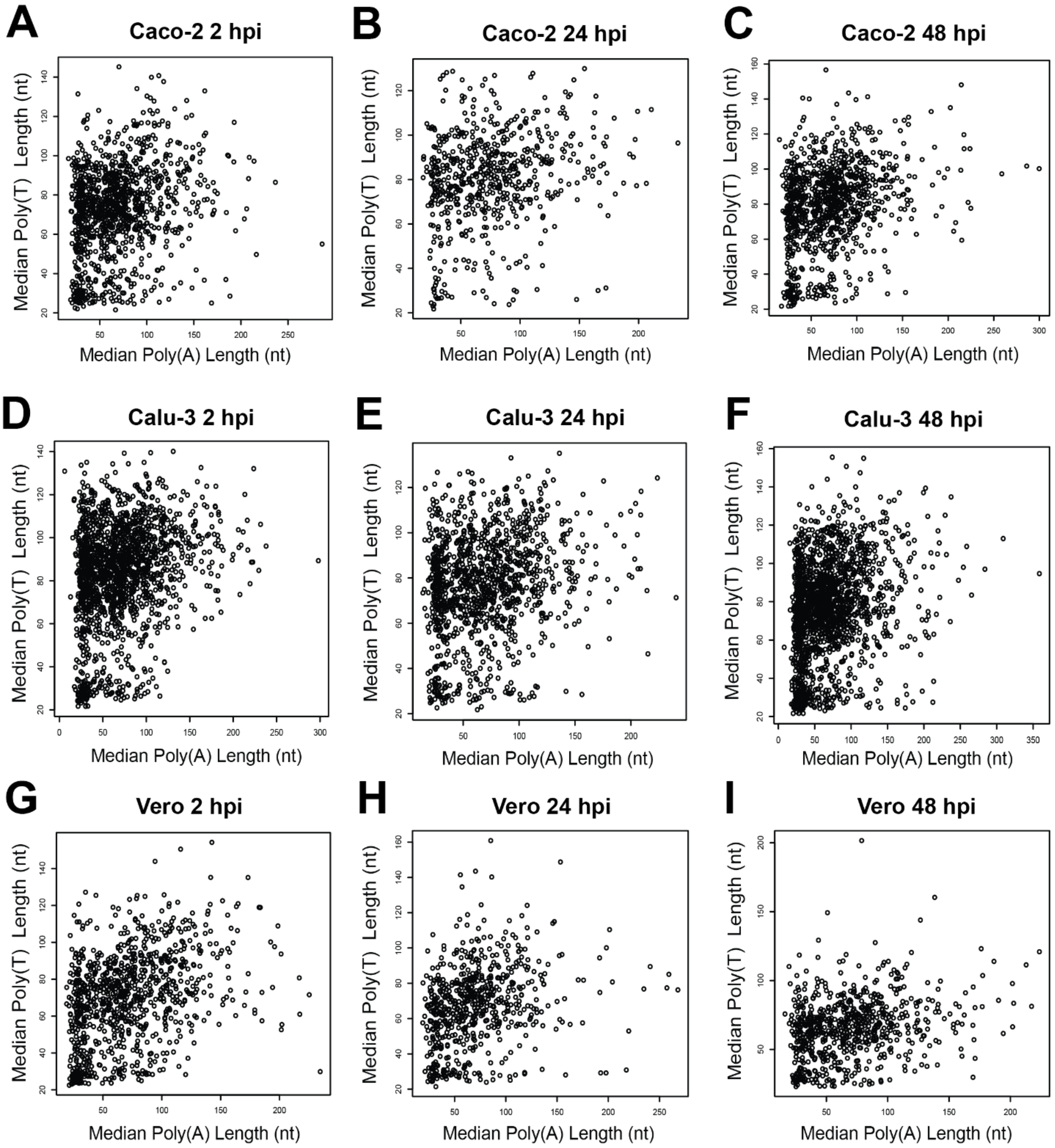


**Figure S3. Correlations between median poly(A/T) lengths from *tailfindr* analyses.** Scatter plots comparing *tailfindr* poly(A) vs poly(T) datasets from Caco-2 **A)** 2 hpi, **B)** 24 hpi, **C)** 48 hpi, Calu-3 **D)** 2 hpi, **E)** 24 hpi, **F)** 48 hpi, Vero **G)** 2 hpi, **H)** 24 hpi**, I)** 48 hpi. Weak correlation between median poly(A) and poly(T) lengths were observed (R < 0.4, exact values along with p-values are indicated in **Table S1**), where each dot represents a gene. Related to **Tables 4 & S1-2**.

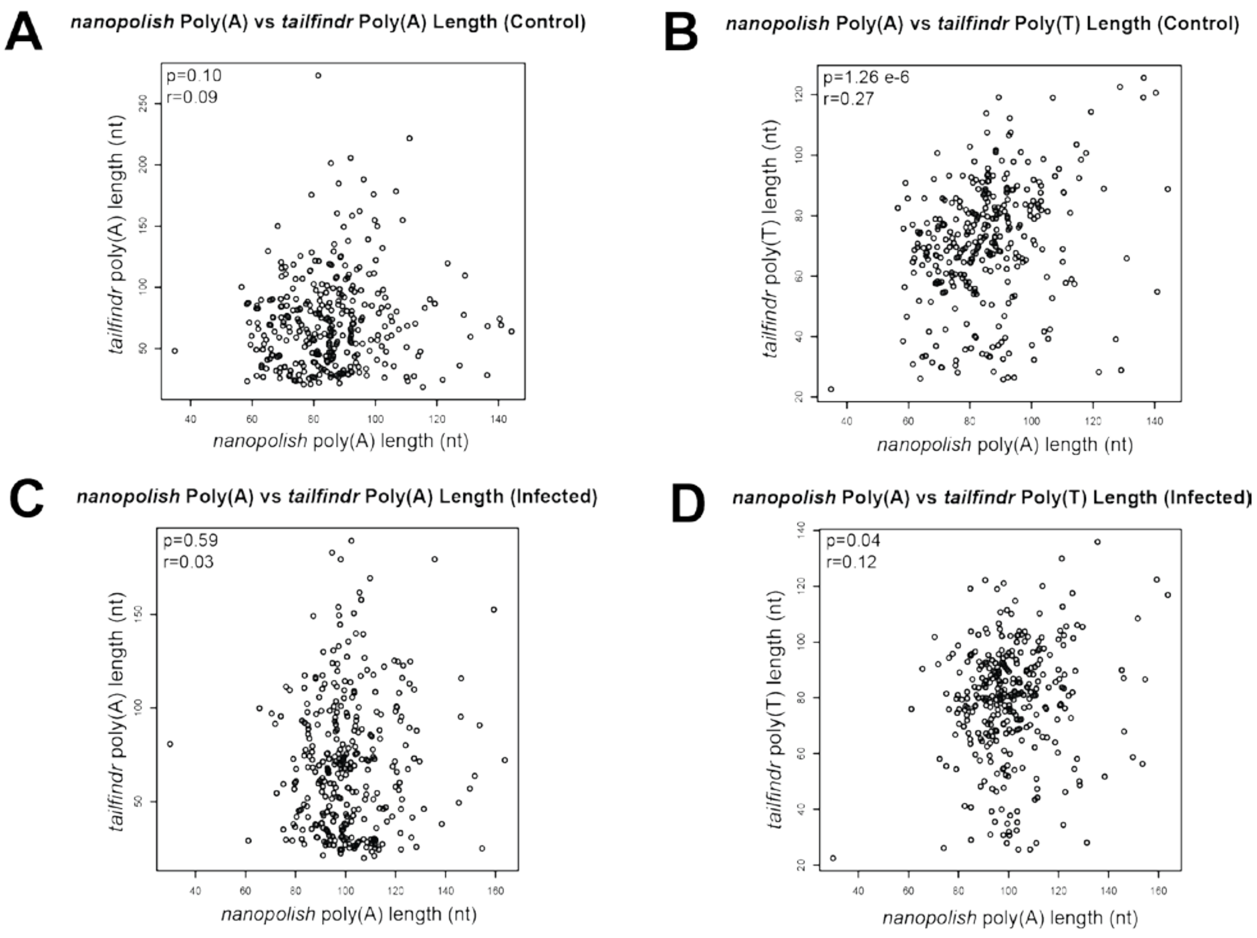


**Figure S4. Correlations between median poly(A) lengths from *nanopolish* and poly(A/T) lengths from *tailfindr* analyses.** Scatter plots comparing **A)** median *nanopolish* poly(A) and *tailfindr* poly(A) lengths in control cells **B)** median *nanopolish* poly(A) and *tailfindr* poly(T) lengths in control cells, **C)** median *nanopolish* poly(A) and *tailfindr* poly(A) lengths in infected cells and **D)** median *nanopolish* poly(A) and *tailfindr* poly(T) lengths in infected cells. Weak significant positive correlations were observed between the *nanopolish* poly(A) and *tailfindr* poly(T) datasets (r = 0.12-0.27, p-value < 0.05, Spearman's correlation test). No significant correlation was observed between *nanopolish* poly(A) and *tailfindr* poly(A) datasets (p-value > 0.05, Spearman's correlation test). Related to **Tables 4 & S2**.

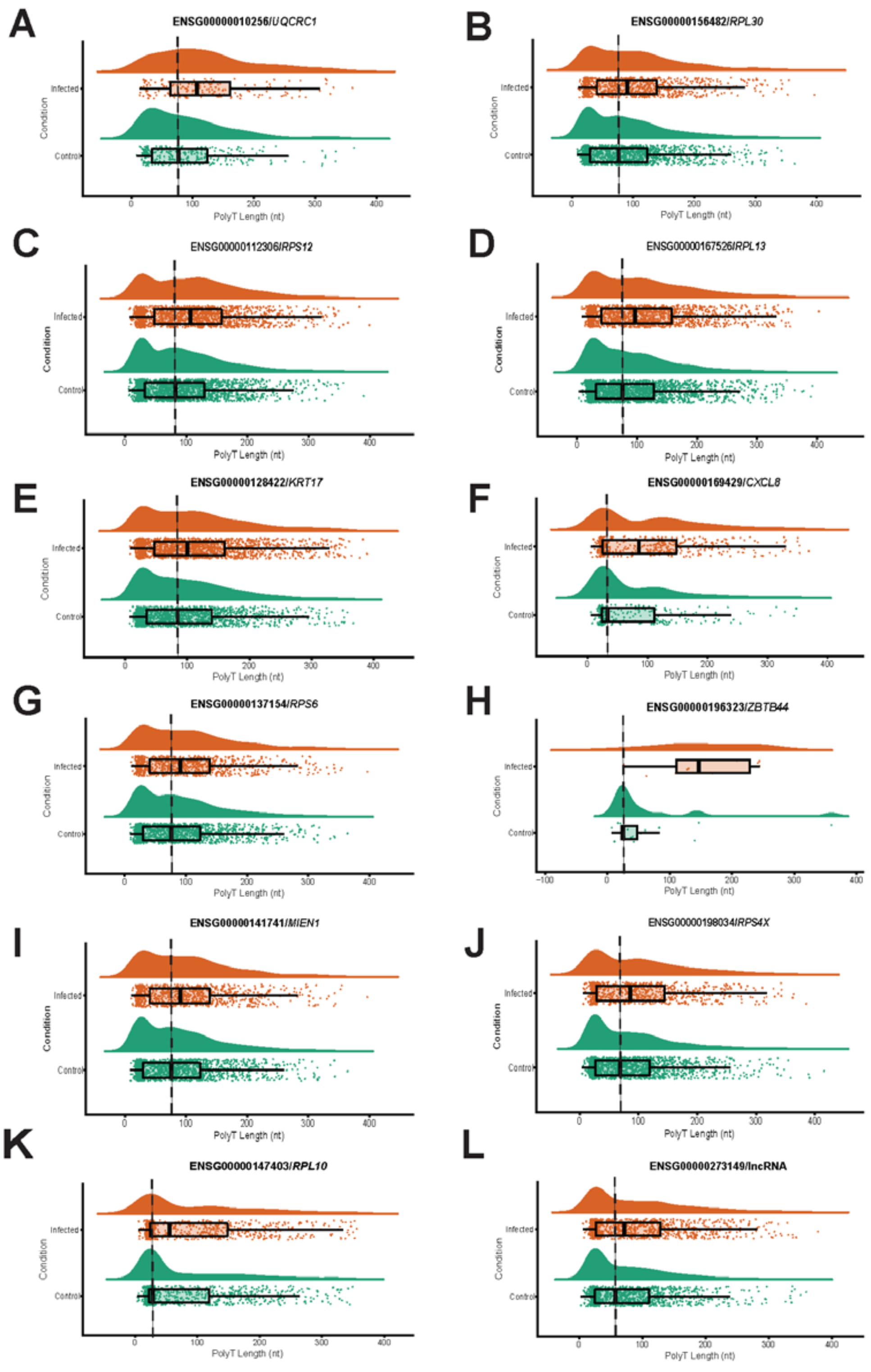


**Figure S5. Raincloud plots of raw, untransformed poly(T) tail lengths in twelve genes with increased poly(A) tail length in the *tailfindr* poly(T) mixed-model analysis with log-transformation.** Genes include **A)** *UQCRC1*, **B)** *RPL30*, **C)** *RPS12*, **D)** *RPL13*, **E)** *KRT17*, **F)** *CXCL8*, **G)** *RPS6*, **H)** *ZBTB44*, **I)** *MIEN1*, **J)** *RPS4X*, **K)** *RPL10* and **L)** a lncRNA. Related to **Figure 4**.

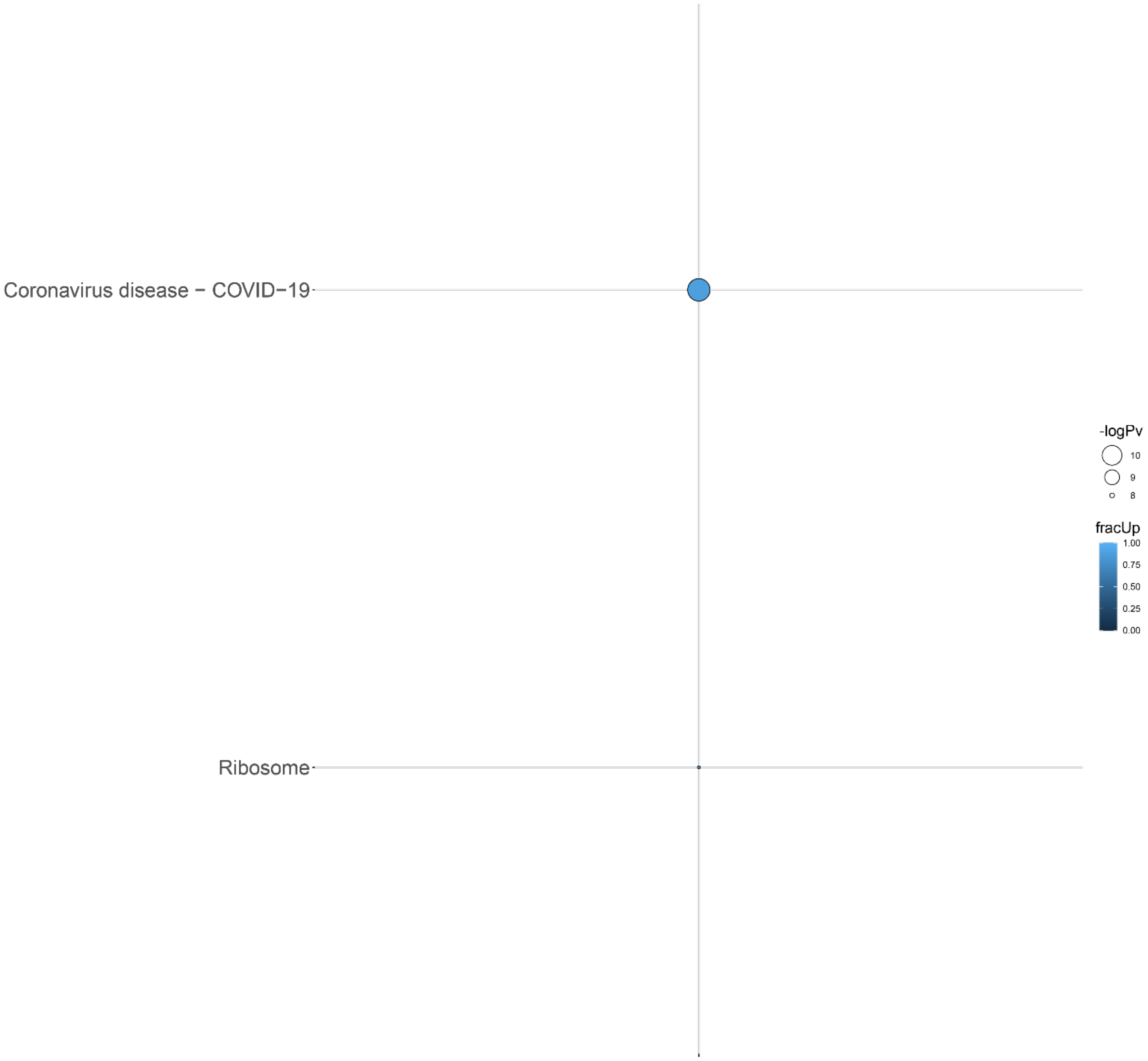


**Figure S6. KEGG pathways from genes with differential poly(A) tail length in *tailfindr* poly(T) mixed-model analyses from the Calu-3 48 hpi dataset.** Genes involved in *coronavirus disease* and *ribosome* pathways were increased in poly(A) length after infection. The bubble size and colour indicate the -log10 enrichment p-values and the fraction of genes with increased polyadenylation, respectively. Thresholds of padj < 0.05, enrichment p-value < 0.0001 were used. Only bubbles which meet the thresholds are shown. Related to **Figure 4 & Data S3**.

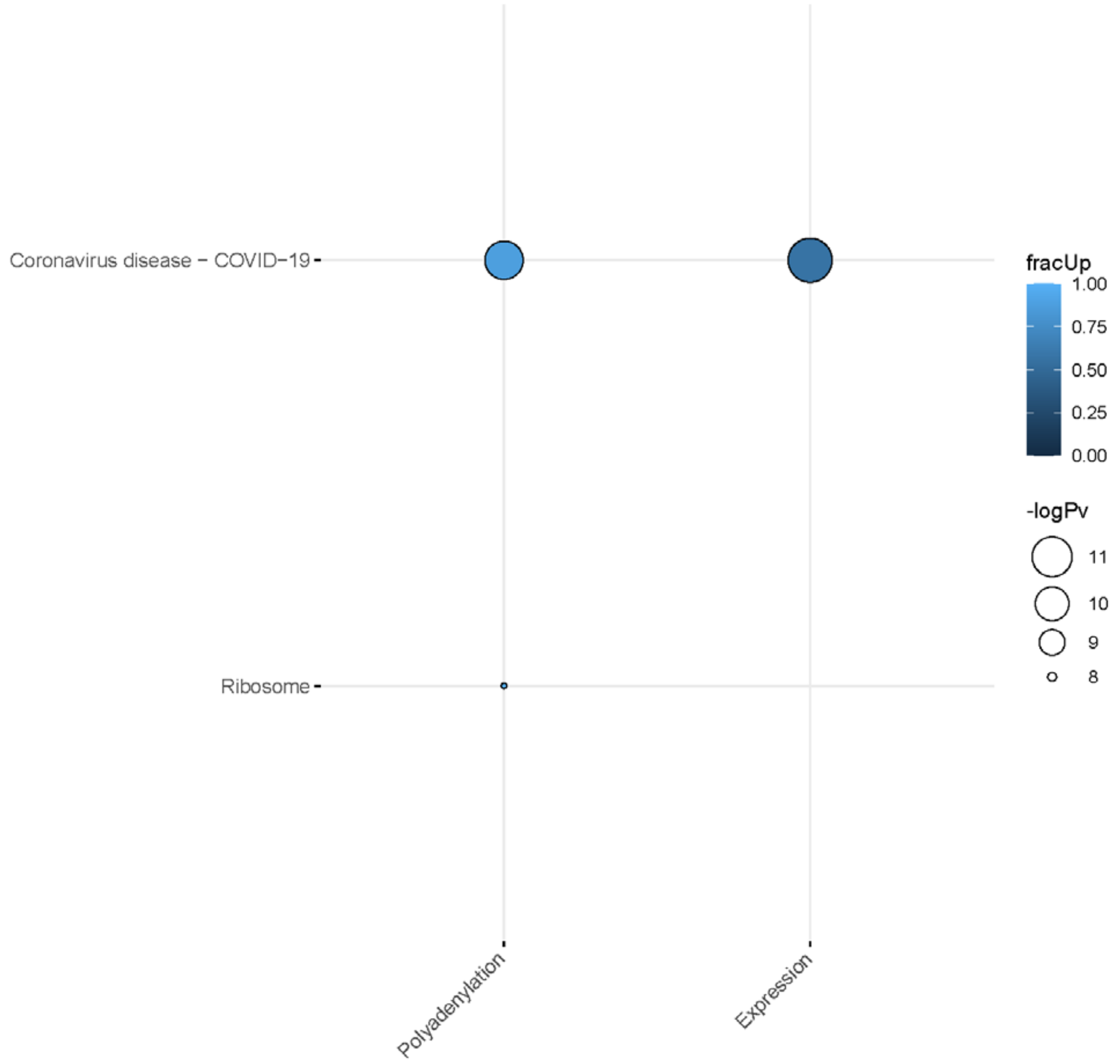


**Figure S7. KEGG pathways of differentially polyadenylated and expressed genes from the Calu-3 48 hpi direct cDNA dataset.** Only the *coronavirus disease* pathway was shared between the two analyses. The plot shows a potential correlation in increased poly(A) tail length and downregulation in gene expression. The bubble size and colour indicate the -log10 enrichment p-values and fraction of upregulated genes/genes with increased polyadenylation, respectively. Thresholds of padj < 0.05, enrichment p-value < 0.0001 were used. Related to **Figure 5 & Data S4**.
